# Multi-omics profiling of cross-resistance between ceftazidime-avibactam and meropenem identifies common and strain-specific mechanisms in *Pseudomonas aeruginosa* clinical isolates

**DOI:** 10.1101/2025.01.14.632673

**Authors:** Bartosz J. Bartmanski, Anja Bösch, Steven Schmitt, Niranjan R. Ireddy, Qun Ren, Jacqueline Findlay, Adrian Egli, Maria Zimmermann-Kogadeeva, Baharak Babouee Flury

## Abstract

*Pseudomonas aeruginosa* is a highly versatile and resilient pathogen that can infect different tissues and rapidly develop resistance to multiple drugs. Ceftazidime-avibactam (CZA) is an antibiotic often used to treat multidrug resistant infections, however, the knowledge on the CZA resistance mechanisms in *P. aeruginosa* is limited. Here we performed laboratory evolution of eight clinical isolates of *P. aeruginosa* exposed to either CZA or meropenem (MEM) in sub-inhibitory concentrations, and used multi-omics profiling to investigate emerging resistance mechanisms. The majority of strains exposed to MEM developed high resistance (83%, 20/24 strains from eight clinical isolates), with only 17% (4/24) acquiring cross-resistance to CZA. The rate of resistance evolution to CZA was substantially lower (21%, 5/24), while 38% (9/24) acquired cross-resistance to MEM. Whole-genome sequencing revealed strain heterogeneity and different evolutionary paths, with three genes mutated in three or more strains: *dacB* in CZA-treated strains and *oprD* and *ftsI* in MEM-treated strains. Transcriptomic and proteomic analysis underlined heterogeneous strain response to antibiotic treatment with few commonly regulated genes and proteins. To identify genes potentially associated with antibiotic resistance, we built a machine learning model that could separate CZA- and MEM-resistant from sensitive strains based on gene expression and protein abundances. To test some of the identified associations, we performed CRISPR/Cas9 genome editing that demonstrated that mutations in *dacB, ampD,* and to a lesser extent in *mexR* directly affected CZA resistance. Overall, this study provides novel insights into the strain-specific molecular mechanisms regulating CZA resistance in *Pseudomonas aeruginosa*.

**Importance:** *Pseudomonas aeruginosa* is one of the most difficult-to-treat pathogens in the hospital, which often acquires resistance to multiple antibiotics. Ceftazidime-avibactam (CZA) is an essential antibiotic used to treat multidrug resistant infections, but its resistance mechanisms are not well understood. Here we investigated the evolution of resistance to CZA and meropenem (MEM) in eight clinical bacterial isolates from patients’ blood, urine, and sputum. While the rate of resistance evolution to MEM was higher than to CZA, MEM-resistant strains rarely acquired cross-resistance towards CZA. To identify changes at the genome, transcriptome and proteome levels during antibiotic exposure, we performed multi-omics profiling of the evolved strains, and confirmed the effect of several genes on antibiotic resistance with genetic engineering. Altogether, our study provides insights into the molecular response of *P. aeruginosa* to CZA and MEM and informs therapeutic interventions, suggesting that CZA could still be effective for patients infected with MEM-resistant pathogens.

## Introduction

*Pseudomonas aeruginosa* is an opportunistic pathogen causing serious infections in hospitalized patients, including chronic respiratory infections in patients with cystic fibrosis (1). This pathogen can adapt to diverse ecological niches (2,3), and is considered a difficult-to-treat pathogen because of its versatile antibiotic resistance mechanisms, hypermutable strains, and antibiotic tolerant cells (4–7).

Given a high prevalence of multidrug resistance (MDR) in *P. aeruginosa*, appropriate antibiotic selection is essential. Acute, life-threatening infections must be treated immediately, while antibiotic susceptibility testing results of clinical isolates often takes up to three days to obtain (8). Empiric treatment often includes beta-lactam antibiotics, such as piperacillin-tazobactam, broad spectrum cephalosporins or carbapenems such as meropenem (MEM) (9). Despite their notable efficacy against *P. aeruginosa* (8,10), multiple resistance mechanisms against these antibiotics have emerged (11–14). Ceftazidime-avibactam (CZA), which combines ceftazidime with a non-β-lactam β-lactamase inhibitor, became a promising treatment option with a broad spectrum activity against MDR pathogens (15,16). The introduction of CZA has not been without setbacks. Soon after its release, resistance was described in *Klebsiella pneumonia*e isolates (17,18). Resistance to CZA has also been described *in vitro* in *P. aeruginosa* that were incubated in the presence of increasing concentrations of CZA. All resistant variants exhibited mutations in their *ampC,* mainly in the Omega-loop region (19). In addition, cross-resistance to CZA was revealed in *P. aeruginosa* with single nucleotide polymorphisms (SNPs) in *ampC* and *ampR*, which led to ceftolozane-tazobactam resistance (20,21).

Multi-omics, including transcriptomics or proteomics, are increasingly used to study antibiotic resistance mechanisms across molecular layers (22–24). Complex antibiotic responses reported in *P. aeruginosa* and other pathogens range from changes in energy metabolism, ribosomal activity, and DNA metabolism, to peptidoglycan biosynthesis, and stress response (25,26). While CZA responses include de-repression of cephalosporinase AmpC, efflux pump upregulation, porin-loss/modification, and deficiencies in DNA mismatch repair (27), the exact resistance mechanisms remain unclear (28–30). Another important aspect of managing *P. aeruginosa* infections is the understanding of cross-resistance evolution, particularly in cases previously treated with meropenem (MEM). This information is essential for rapid clinical decision-making, but relevant data on this topic is currently lacking.

This study aims to elucidate resistance and cross-resistance mechanisms to MEM and CZA in *P. aeruginosa* through *in vitro* evolution experiments, multi-omics profiling and genome engineering, offering valuable insights for physicians selecting empirical treatments for *P. aeruginosa* infections pre-treated with meropenem. Additionally, it highlights *P. aeruginosa* strategies for MEM and CZA adaptation and resistance, which could guide the development of new antibiotics and combination therapies.

## Results

### Experimental evolution of antimicrobial resistance to MEM and CZA in clinical isolates

To investigate the evolution of resistance and cross-resistance to MEM and CZA in *P. aeruginosa*, we collected eight clinical isolates from three different infection locations (blood, sputum, and urine), confirmed their sensitivity to the drugs (Supplementary Table 1), and exposed them to sub-inhibitory concentrations of either drug for 18 days in triplicates (Figure 1a). We observed that among the strains exposed to sub-inhibitory levels of CZA 21% (5/24) became resistant to CZA, while 38% (9/24) demonstrated the development of cross-resistance to MEM in accordance with the European Committee on Antimicrobial Susceptibility Testing (EUCAST) resistance breakpoint guidelines of MIC>8 mg/L (31) (Figure 1b, Supplementary Figure 1a). In contrast, most strains (20/24, 83%) subjected to sub-inhibitory concentrations of MEM acquired MEM resistance, with only a minority of 17% (4/24) concurrently displaying cross-resistance to CZA (Figure 1c, Supplementary Table 2). This observation also holds true if we consider a subset of five isolates which had a low initial MEM MIC (MIC<=1 mg/L) (Supplementary Figure 1b). The increase of MIC was much higher for MEM compared to CZA (median increase of 32-fold and 5.3-fold versus 2.7-fold and 3-fold upon exposure to MEM and CZA, correspondingly, Supplementary Table 2).

**Figure 1.**
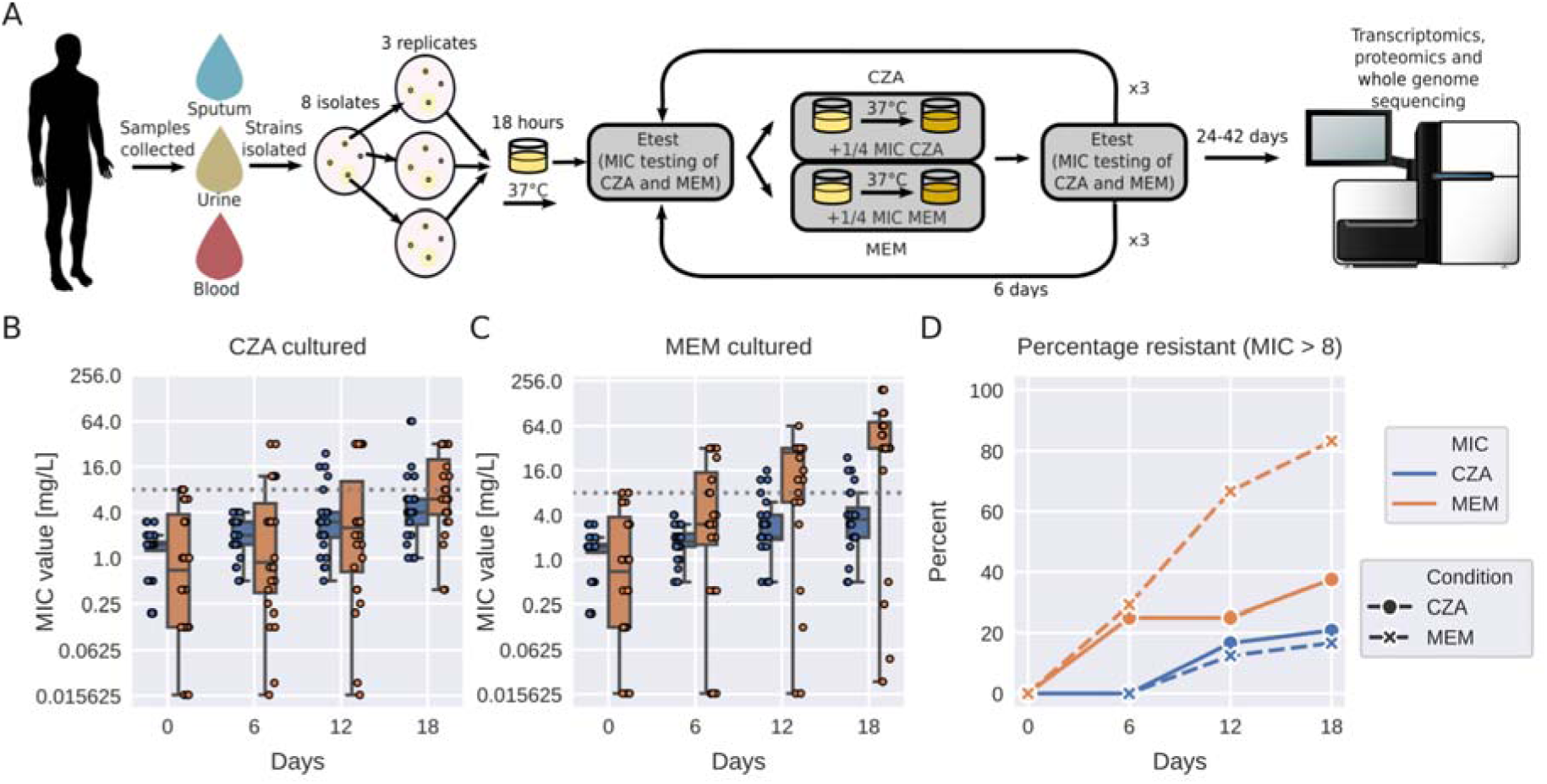
Experimental evolution of antimicrobial resistance to MEM and CZA in clinical isolates. (A) Experimental design. (B) Resistance evolution to MEM and CZA in strains exposed to CZA. (C) Resistance evolution to MEM and CZA in strains exposed to MEM. (D) Percentage of MEM- and CZA-resistant strains (MIC>8 mg/L) among strains exposed to either MEM or CZA. In (B) and (C), gray horizontal dotted line corresponds to the MIC=8 mg/L threshold. Boxplots represent median and interquartile range (between the first and the third quartile), while the whiskers extend from the box to the farthest data point lying within 1.5x the interquartile range from the box for n=24 points (n=8 strains in triplicates). MIC - minimal inhibitory concentration. In (A), genomesequencer-2 icon by DBCLS https://togotv.dbcls.jp/en/pics.html is licensed under CC-BY 4.0 Unported https://creativecommons.org/licenses/by/4.0/. petri-dish-with-colony-lightyellow icon by Servier https://smart.servier.com/ is licensed under CC-BY 3.0 Unported https://creativecommons.org/licenses/by/3.0/.

### MEM and CZA induce distinct molecular responses across multi-omics layers

Given that only a minority, constituting 21% of the strains exposed to CZA, had exceeded the clinically defined resistance breakpoint of MIC>8 mg/L (31) after 18 days, we extended the evolutionary experiment for a longer period of time to capture the molecular mechanisms underlying the development of resistance. Specifically, one representative strain evolved from each of the parental isolates was subjected to sub-inhibitory concentrations of the respective antibiotics for a maximum of 42 days of continuous passaging. For technical reasons, we were able to collect the data from six strains evolved in CZA and five strains evolved in MEM out of eight parental strains. After this prolonged period, four out of six strains acquired CZA resistance (MIC>8 mg/L), with two of them being cross-resistant to MEM (MIC>8 mg/L), and one having the intermediate resistance. One of the CZA-sensitive strains also had intermediate MEM resistance. While all five MEM-exposed strains acquired high resistance (MIC>96 mg/L), only two of them acquired CZA cross-resistance (Figure 2a, Supplementary Table 3). Notably, during exposure to MEM and CZA, strains also evolved resistance to other beta-lactams, cephalosporins and other antibiotics (Supplementary Table 4).

For all the parent and the evolved strains, we performed whole genome sequencing, and transcriptomics and proteomics analysis with three technical replicates to investigate acquired mutations and changes in gene expression and protein abundance upon antibiotic treatment.

Whole genome sequencing revealed that the six strains carried five sequence types (ST) and were genetically diverse, representing different clades of *P. aeruginosa* phylogenetic tree (Supplementary Figures 2 and 3). Most evolved strains accumulated a similar number of mutations when exposed to MEM or CZA (between three and twenty nine), while one of the strains (1_B_) was a hypermutable strain, accumulating more than 150 mutations (Figure 2b, Supplementary Table 5). Several genes acquired mutations independently in several strains or in the same strain exposed to either of the drugs. The most frequently mutated genes were penicillin binding proteins *ftsI* (in two strains exposed to CZA and four strains exposed to MEM) and *dacB* (in four strains exposed to CZA). These were followed by outer membrane protein *oprD* (in three strains exposed to MEM), two-component system sensor *phoQ* (in two strains exposed to MEM and one strain exposed to CZA), and multi-drug efflux pump *mexB* and its regulator *mexR* (in one strain exposed to CZA and two strains exposed to MEM) (Supplementary Table 5). However, none of the mutations could be unambiguously attributed to the levels of acquired CZA or MEM resistance.

Exposure to MEM or CZA induced large changes in gene expression (up to 1390 differentially expressed genes across strains, abs(log2(fold change)) > 1, FDR-adjusted p-value < 0.05) and protein abundance (up to 888 differentially abundant proteins, abs(log2(fold change))>1, FDR-adjusted p-value < 0.05) (Figure 2c, Supplementary Tables 6 and 7, Supplementary Figures 3 and 4). While gene and protein responses were distinct between the two drugs (up to 1329 differentially expressed genes and 888 differentially abundant proteins per strain between MEM and CZA conditions), gene ontology enrichment analysis revealed several common processes affected by either of the antibiotic treatments. These included membrane transport, protein secretion and cellular localization. Furthermore, exposure to CZA and MEM affected amino acid, lipid and nonribosomal peptide metabolic processes in different strains (Figure 2d, Supplementary Table 8).

While we observed some common responses to MEM and CZA exposure between strains, there was no process that could be unambiguously attributed to antimicrobial resistance. For example, while several genes involved in protein transport, localization and secretion were regulated in CZA-resistant strains 1_B_ and 2_C_ upon CZA exposure, they were not enriched among genes regulated by the two other resistant strains 4_C_ and 5_B_. The response to CZA of the highly CZA-resistant strain 5_B_ was unique to this strain and characterised mainly by the changes in biosynthetic and metabolic processes. Therefore, next we set out to compare molecular changes across strains in more detail to identify potential antibiotic-resistance features.

**Figure 2.**
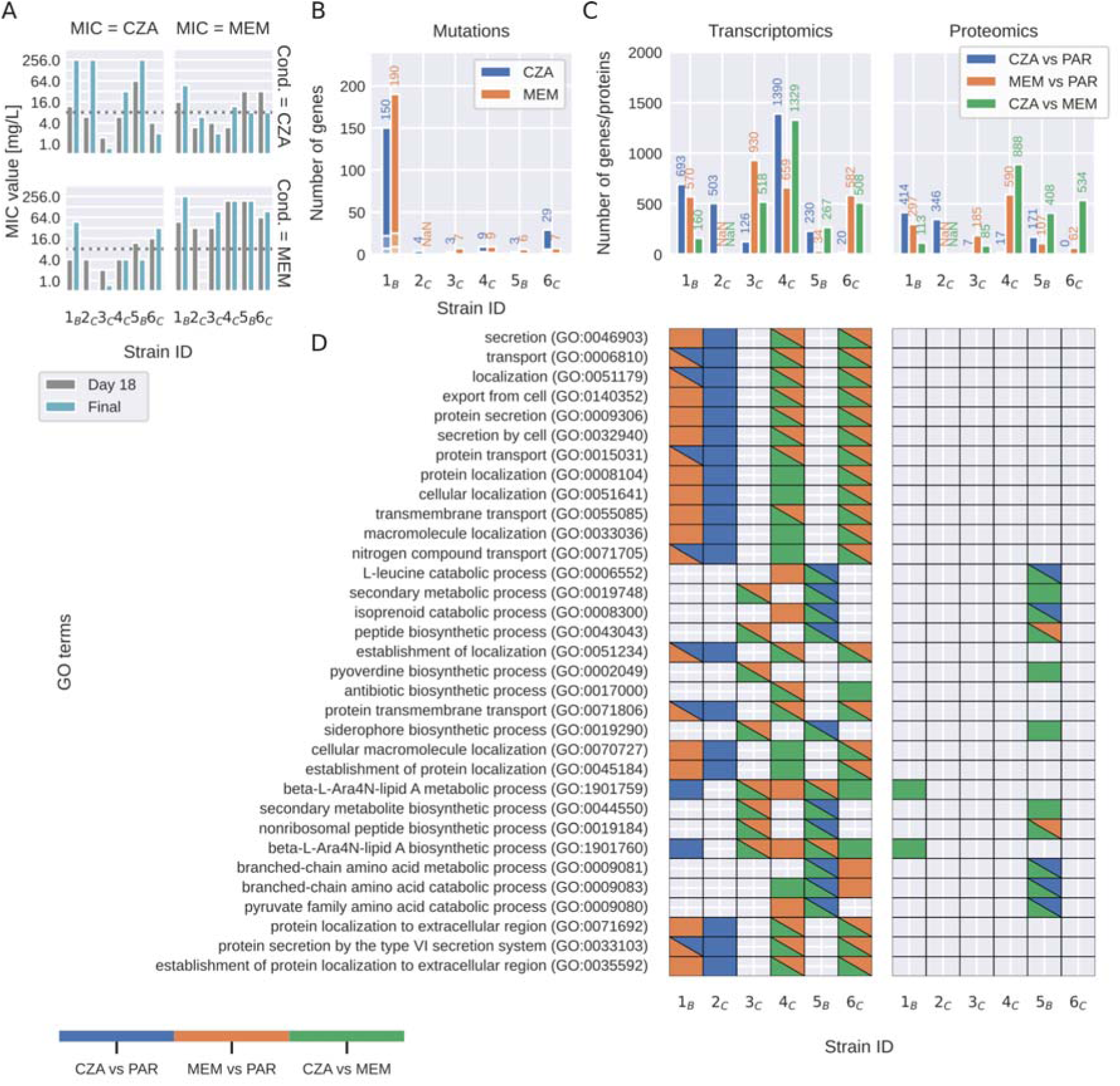
Molecular responses to MEM and CZA exposure across clinical isolates. (A) MIC values for strains exposed to either CZA or MEM after the initial 18 days of passaging and after 24-42 days of passaging. (B) Number of mutations induced by MEM or CZA exposure (depicts the sum of all types of detected mutations, such as SNP, insertion, deletion and complex rearrangements). (C) Number of differentially expressed genes and differentially abundant proteins induced by MEM or CZA exposure compared to parent strains (PAR) (log2(fold change)>=1, FDR<=0.05 determined using Wald test and unpaired two-sided moderated t-test for gene expression and protein abundance, respectively, and adjusted with Benjamini–Hochberg procedure). (D) GO enrichment analysis of differentially abundant genes and proteins. Color highlights significantly enriched GO terms (FDR<=0.05). For visualization purposes, only GO terms that pass a significance threshold of FDR<=0.001 in at least two strains for transcriptomics or in at least one strain for proteomics are shown.

### Molecular signatures of evolved strains are largely specific to the parental strain

To compare molecular signatures between strains and conditions we performed dimensionality reduction of the normalised transcriptomics and proteomics profiles with principal component analysis (PCA). This analysis revealed that the evolved strains exposed to either MEM or CZA grouped together with the parental strain, underlining that the parental strain determined the global transcriptional and protein abundance profiles of the evolved strains more than the antibiotic exposure (Figure 3a). We note that due to insufficient sample quality, we had to exclude the MEM-evolved 2_C_ strain from our analysis.

Indeed, while MEM or CZA treatment induced large transcriptional (1895 or 2178 out of 5563 detected genes) and protein (984 or 809 out of 4028 detected proteins) changes across strains, most of these changes were strain-specific (Figure 3b, Supplementary tables 6, 7 and 9). MEM induced more common transcriptional and protein changes across strains than CZA. About 41% of proteins changing upon MEM exposure (401 out of 984) and 40% of proteins changing upon CZA exposure (320 out of 809) were also changing on the transcript levels. They were mainly enriched in the nucleic acid metabolic processes, phenazine biosynthesis, and nitrogen compound transport (GO:0090304, GO:0002047 and GO:0071705) in MEM, and pyruvate family amino acid and branched-chain amino acid catabolic processes in CZA (GO:0009080 and GO:0009083, Supplementary Table 8). The only genes significantly upregulated by four strains upon MEM treatment were *mexA* and *mexB,* and the corresponding proteins were upregulated as well (Figure 3b, Supplementary Table 9). While no proteins were differentially abundant in more than three strains cultured in CZA, there were seven genes that were significantly up- or down-regulated in five strains cultured in CZA: porin B *oprB* (PA3186), beta-lactamase *ampC* (PA4110) and ribosomal protein S3AE (PA4111), as well as four genes belonging to an operon of ABC transporter genes PA3187-PA3190 (Supplementary Table 9). Since these genes were differentially expressed in both CZA-resistant and sensitive strains, they are likely not unambiguously determining resistance, but reflecting the stress response and bacterial attempts to extrude the antibiotic. To put the findings observed for the six clinical isolates into a broader perspective, we leveraged publicly available datasets on *P. aeruginosa* PAO1 strain exposed to MEM (32,33), as well as transcriptomic profiles of large clinical isolate collections tested for MEM or CZA resistance (34,35). While *mexA* was highly expressed in CZA-resistant strains and not differentially expressed by MEM-treated PAO1 strain, *ampC* and PA4111 were highly expressed both in MEM-resistant strains and in PAO1 treated with MEM. Three more genes out of the nine genes discussed above were also upregulated in PAO1 strain in response to MEM (PA3186-88, Supplementary Table 10), supporting that they are rather markers of MEM-response but not unambiguous resistance determinants. *P. aeruginosa* PAO1 response to MEM shared up to 40% of differentially expressed genes with the clinical isolates, providing an additional reference for general drug response of an antibiotic-sensitive strain (Supplementary Figure 6, Supplementary Table 11).

**Figure 3.**
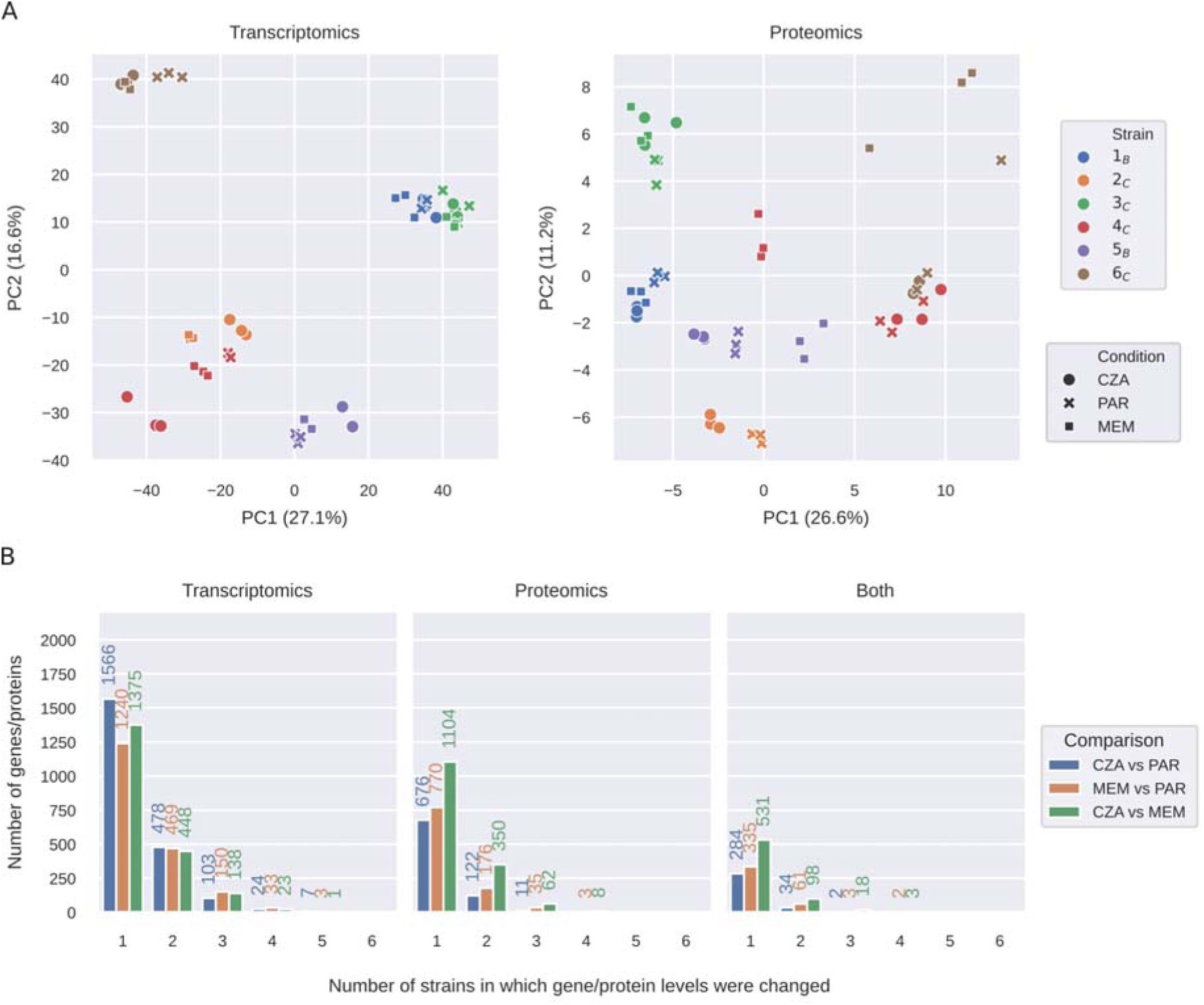
Molecular signatures of evolved strains are largely specific to the parental strain. (A) PCA plots of normalised transcriptomics and proteomics data for the parent strains and strains evolved in MEM or CZA. (B) Number of differentially expressed transcripts, differentially abundant proteins, or both, shared between each number of strains (log2(fold change)>=1, FDR<=0.05 determined using Wald test and unpaired two-sided t-test for gene expression and protein abundance, respectively, and adjusted with Benjamini–Hochberg procedure). PAR refers to parental strains.

### A machine learning model identifies genes and proteins predictive of MEM and CZA resistance

To investigate whether MEM- or CZA-resistant strains have distinctive molecular signatures, we used a machine learning method Partial Least Squares-Discriminant Analysis (PLS-DA), which is widely applied for discriminative analysis and classification in high dimensional datasets (36,37). PLS-DA aims to maximise the covariance between the sample class (defined as high resistance to CZA or MEM, MIC>=32 mg/L) and the predictor features (gene expression or protein abundance), and extracts latent features (latent factors) that capture the essential information for discriminating between the two classes. First, we built four PLS-DA models (for each of the two antibiotics and each of the two omics data types) using the full dataset, including parent and CZA- and MEM-evolved strains (51 samples per model). All four models could separate resistant from sensitive strains with high accuracy (accuracy 0.96 and higher) (Figure 4a, Supplementary Table 12). To test the generalizability of the models, we performed a leave-one-strain-out analysis, where each of the models was trained on the data from five strains (42-45 samples) and tested on the data from the sixth strain not included in the training (6-9 samples) (Figure 4b). Reflecting the heterogeneity between strains, resistance prediction accuracy substantially varied depending on which strain was left out (Figure 4c). Both MEM- and CZA-resistance models trained on proteomics data achieved higher accuracy than models trained on transcriptomics data (median accuracies 0.72 and 0.67 compared to 0.67 and 0.36, Supplementary Table 12).

We then took advantage of the latent factor projections calculated by PLS-DA to identify genes and proteins most informative for the prediction of each model. For each of the gene or protein features, we calculated how many leave-one-strain-out models included this feature among the top 100 features sorted by the latent factor projection coefficients (out of 5563 total features for transcriptomics and 1923 total features for proteomics). While most of the top predictive features were model-specific, we identified small subsets of features that were repeatedly picked by five or all six leave-one-strain-out models of each type (Figure 4d, Supplementary Table 12). For MEM resistance prediction, the models based on transcriptomics picked genes involved in membrane functions, while most frequently picked proteins were involved in metabolic processes, specifically carbohydrate derivative, carnitine and dTDP-rhamnose metabolism (Figure 4e, Supplementary Table 12). For CZA resistance prediction, the selected gene features were attributed to aminoglycan and peptidoglycan catabolic processes, while protein features were involved in gene expression, protein folding and cysteinyl-tRNA aminoacylation (Figure 4e, Supplementary Table 12). We then focused on the features selected by five or more MEM- or CZA-predicting models (40 and 24 features correspondingly, Figure 4d), and filtered them further by calculating whether they were significantly differentially expressed between CZA-sensitive and resistant strains (FDR<=0.01, Supplementary Table 13). This resulted in 9 features for MEM-resistance and 12 features for CZA-resistance (Supplementary Figures 5 and 6, Supplementary Table 14). Among the 12 CZA-resistance features, three genes were reported to affect resistance to different antibiotics in a genome-wide transposon mutant study (30) (Supplementary Table 14). Two of these genes are associated with alginate biosynthesis (PA0460 and PA0764 (*mucB*)), while the third one, glutathione hydrolase proenzyme *ggt* (PA1338) was also reported to be downregulated in a gentamicin-resistant clinical isolate (39). Alginate and other exopolysaccharides are known to affect biofilm formation, pathogenicity and antibiotic resistance (40–43). The other genes from the list are involved in cell wall biosynthesis (PA4421 (*mraZ*)) or metabolic functions (PA2197, PA3945, PA4847 (*accB*) PA3082 (*gbt*) PA5556 (*atpA*)) and have been associated with virulence, antibiotic response or resistance (44–48), while the upregulated selenide, water dikinase *selD* (PA1642) was proposed as a drug target in *Klebsiella* (49) (Supplementary Table 14). Overall, PLS-DA enabled identification of transcriptomics and proteomics features that are associated with a higher resistance to MEM or CZA, which may include antibiotic resistance mechanisms, as well as stress response to the higher antibiotic levels in the antibiotic-resistant group, or general characteristics of highly pathogenic strains, such as biofilm formation capacities.

**Figure 4.**
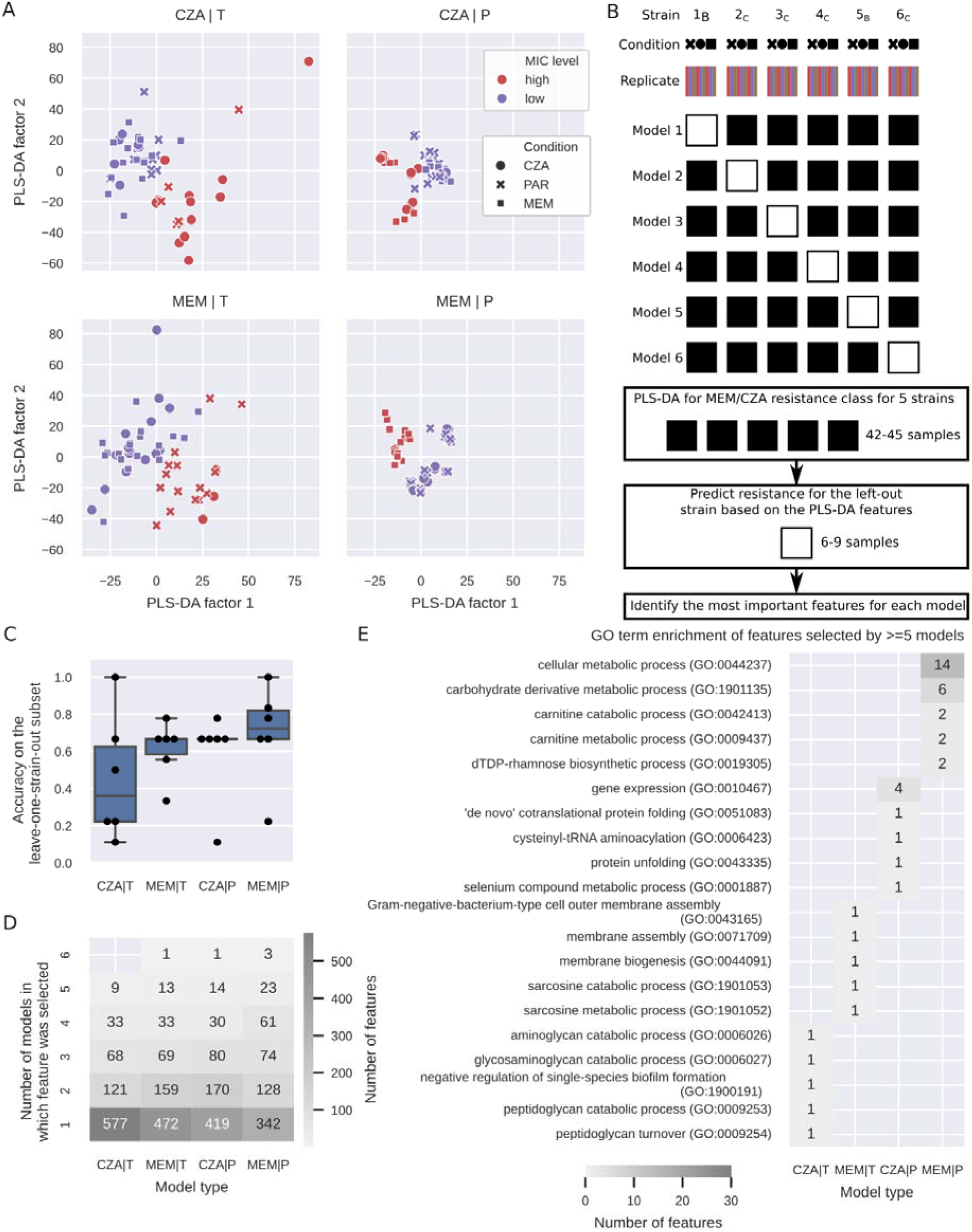
A machine learning model identifies genes and proteins predictive of MEM and CZA resistance. (A) PLS-DA plots for models built for transcriptomics (left) or proteomics (right) data to predict high CZA (top) or MEM (bottom) resistance (threshold at MIC>=32 mg/L). (B) Scheme of the leave-one-strain-out approach to identify genes and proteins predictive of CZA or MEM resistance. (C) Accuracy of the six leave-one-strain-out models on the leave-one-strain-out subset (calculated based on n=6-9 samples for each left out strain) for each of the omics data types and antibiotics. (D) Number of predictive features selected by one to six of the leave-one-strain-out models (only top 100 features per model were considered for comparison). (E) GO enrichment analysis of features selected by >=5 leave-one-strain-out models. The top five GO terms per model are shown sorted by the p-value. Only GO:1901135 passes the FDR<=0.1 threshold.

### Antibiotic exposure-associated features span from membrane and transport processes to energy and lipopolysaccharide metabolism

To summarise antibiotic exposure-associated processes, we combined genes and proteins identified with mutation analysis, differential transcriptomic and proteomic analyses as well as the PLS-DA models. From mutation analysis, we selected genes in which mutation occurred at least two times independently in our assay (19 genes); from differential analysis we selected genes and proteins significantly up- or downregulated (abs(log2(fold change))>=1 in either of the omics, FDR<0.05 in both omics) by at least five strains from transcriptomics, four strains from proteomics and three strains from both (17 genes/proteins). Lastly, from PLS-DA we selected features that were highly important in at least five leave-one-strain-out models and additionally differentially abundant between sensitive and resistant stains (FDR<0.01, 21 genes/proteins, Supplementary Table 14). Since some genes or proteins were selected by multiple of these analyses, there were 56 features selected in total (Supplementary Table 15).

Mutations most frequently occurred in penicillin binding proteins *dacB* (only upon CZA treatment) and *ftsI* (both upon MEM and CZA treatment), while the expression of these genes was unchanged or decreased (Figure 5, Supplementary Table 15). The most pronounced upregulation both on the transcript and the protein levels was observed in *mexAB-oprM* efflux system in the majority of strains exposed to MEM and two of the three strains which acquired high resistance to CZA, however, this upregulation was not coinciding with cross-resistance. Mutations were also detected in the *mexB* gene upon both CZA and MEM exposure, in the *nalD* gene that was reported to contribute to the increase of *mexAB-oprM* expression in *P. aeruginosa* (50), and in the *mexR* gene, recognized as a repressor gene of the *mexAB-oprM* Multidrug Efflux Operon (51) (Figure 5, Supplementary Table 15). The expected antibiotic resistance determinant beta-lactamase *ampC* was upregulated in the majority of the strains exposed to CZA, however, it did not exclusively determine CZA resistance or cross-resistance, since both CZA-sensitive and resistant strains had high *ampC* expression levels according to the transcriptomic profiling. Antibiotic exposure-associated genes and proteins belonging to metabolism and energy production category were selected either by differential analysis or PLS-DA and were mainly downregulated, indicating that antibiotic exposure impaired growth and metabolism of resistant strains more than sensitive strains, likely due to the differences in the administered antibiotic doses.

The two categories of genes/proteins identified by all three types of analysis were membrane proteins and outer membrane channels and polysaccharide and lipopolysaccharide metabolism (Figure 5, Supplementary Tables 15). Mutations occurring within *oprD* were uniquely observed in strains exposed to MEM, but not coinciding with cross-resistance to CZA. The expression of *oprD* as well as the outer membrane protein *oprE* were generally downregulated (Figure 5, Supplementary Table 15). Genes involved in lipopolysaccharide metabolism were mutated mainly upon MEM exposure (the *phoP*-*phoQ* two-component regulatory system, murein peptide ligase *mpl*, and cell division protein *ftsZ*), while the expression of these and other genes in this category was generally increasing both upon MEM and CZA exposure.

To summarise, three types of analysis point towards mutations and expression changes in penicillin binding proteins, efflux pumps membrane proteins, as well as lipopolysaccharide metabolism genes as the most generally associated with antibiotic exposure, while highlighting drug-specific responses, such as mutations in *dacB* upon CZA exposure or *oprD* upon MEM exposure. These affected processes likely reflect not only antimicrobial resistance strategies, but also bacterial stress response and general virulence factors.

**Figure 5.**
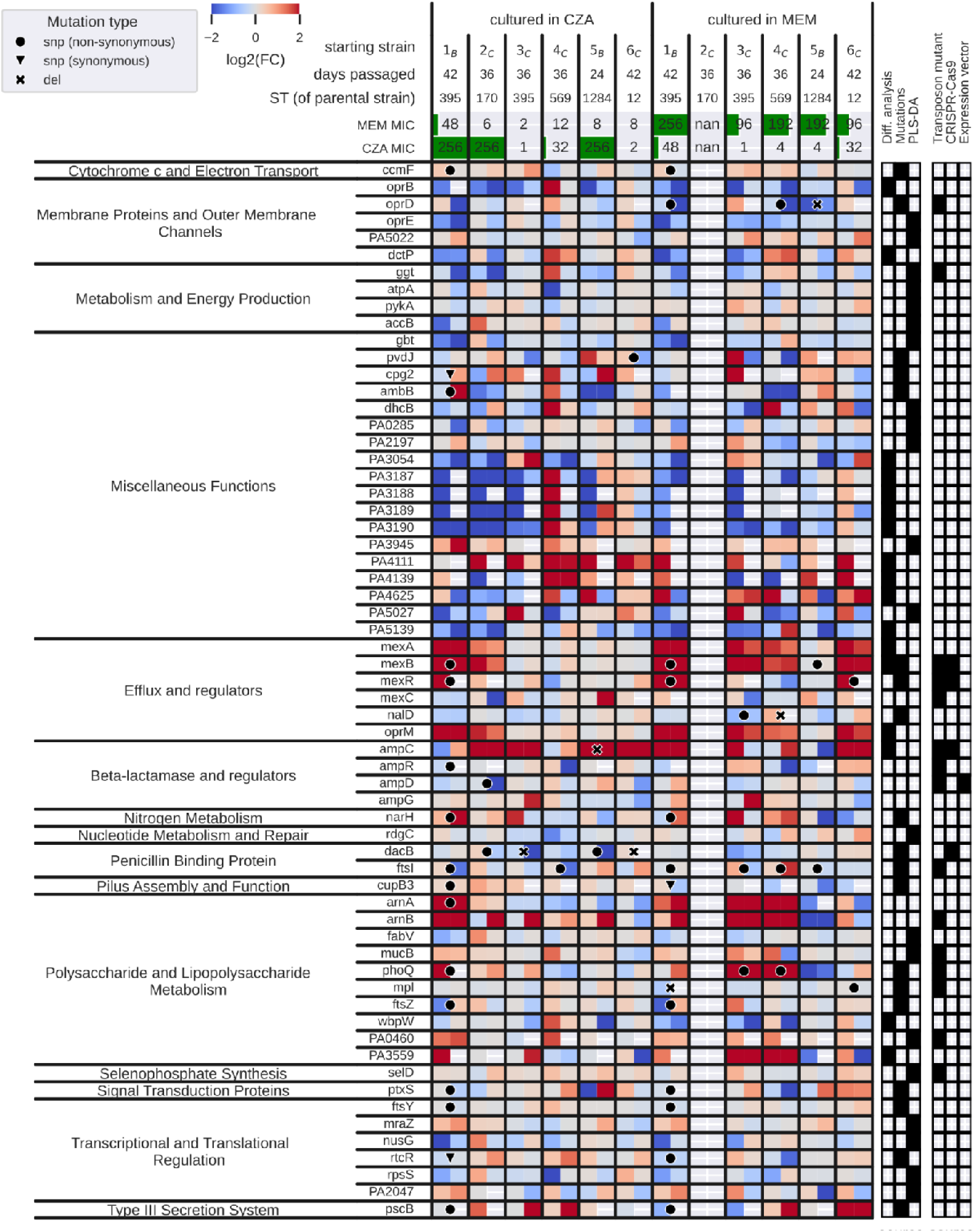
Overview of the antibiotic exposure-associated features identified by different types of analyses. The heatmap depicts transcriptomics log fold changes on the left and proteomics log fold changes on the right within each box for a given strain and gene. Red; up-regulated, blue; down-regulated. Mutations found for each gene in a given strain are depicted by a circle (non-synonymous SNP), a triangle (synonymous SNP) or a cross (deletion). The top two rows show the MIC values for CZA and MEM after the specified number of days. The panels on the right indicate the type of analysis through which each gene was identified (mutations, differential analysis, PLS-DA), and whether the gene was followed up in a CRISPR-Cas9, a transposon mutagenesis assay, or an expression vector cloning assay. Note that due to technical reasons we removed MEM-exposed 2_C_ strain from all the analyses.

### CRISPR-Cas9 genome editing reveals that dacB mutation has a strong effect on CZA resistance

To test whether specific identified mutations can affect CZA or MEM resistance in *P. aeruginosa*, we set out to perform CRISPR-Cas9 genome editing of the *P. aeruginosa* mPAO1 strain using gene sequence templates from the highly resistant strains. Specifically, we focussed on several of the most frequently mutated genes *dacB*, *mexB* and *mexR*, the heavily upregulated *ampC* and its regulator *ampD* (Figure 5).

The mutations in *mexR* and *mexB* were introduced from strain 1_B_, which exhibited high levels of resistance to both CZA and MEM. The introduction of the L13P (T38C) mutation in *mexR*, which is accompanied by a 20-fold increase in its expression upon MEM treatment of 1_B_ (Supplementary Table 6), resulted in a 3-fold rise in MEM MIC and 2-fold rise in CZA MIC respectively. Introducing the R620C (C1858T) mutation in *mexB* led to a 1.5-fold increase in CZA MIC without affecting MEM MIC (Table 1).

For the *ampC* deletion introduced based on the template from isolate 5_B_, we also observed a moderate increase of 1.5 fold in CZA resistance and no effect on MEM resistance (Table 1), likely explained by the fact that this deletion was upstream of the avibactam binding site.

The strongest effect on CZA MIC was observed by introducing the *dacB* mutation acquired by strain 2_C_ exposed to CZA (Table 1): the CZA MIC increased four fold, while no discernible impact on MEM resistance was detected. In contrast, introducing another *dacB* mutation from CZA-resistant strain 5_B_ did not result in any changes in the MIC of either CZA or MEM (Table 1).

Overall, our CRISPR-Cas9 assays could confirm the effects on CZA resistance of *dacB* and *mexR*, although the effects were much less pronounced than the resistance patterns of the evolved strains. Along with CRISPR-Cas9 assays, we also performed MIC assays with a subset of transposon insertion mutants in the *P. aeruginosa* mPAO1 mutant library (n=15 mutants, Figure 5, Supplementary Table 16). From the tested transposon mutants, we observed substantial differences to the wildtype CZA MIC in two out of fifteen: 25% reduction in MIC in *mexB* mutant and 3-fold increase in MIC in *mexR* mutant (Supplementary Figure 9, Supplementary Table 16). These results underline the composite nature of antibiotic resistance mechanisms, as genetic perturbations of single genes cannot fully recapitulate the resistance phenotypes of the evolved strains.

**Table 1.**
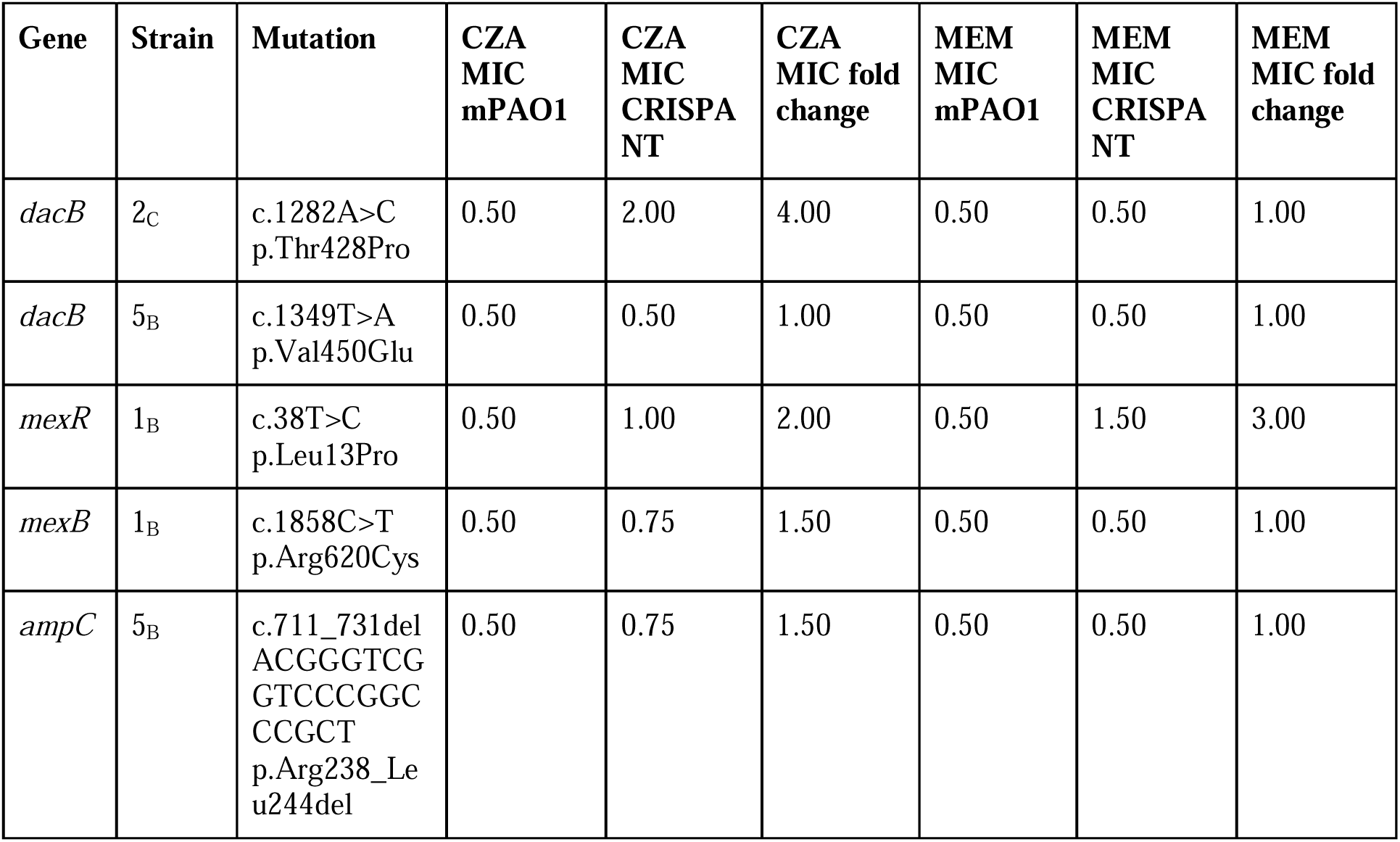
Results of CZA and MEM resistance assays of the CRISPR/Cas9 edited mutants of P. aeruginosa mPAO1.

### Introduction of the wild-type ampD gene reversed CZA resistance in the mutant strain

As CRISPR-Cas9 genome editing was unsuccessful for the *ampD* mutation despite several attempts, we opted for an alternative approach. We cloned the wild-type *ampD* allele from the sensitive 2_C_ parent strain (CZA MIC=2 mg/L) into the pUCP24 vector and introduced it into the evolved 2_C_ strain (CZA MIC=128 mg/L) via transformation assays using *Escherichia coli* TOP10. Complementation with the parent *ampD* gene led to a 64-fold decrease of resistance in the mutant strain (CZA MIC=2 mg/L, Table 2), confirming that *ampD* mutation was the leading cause of acquired CZA resistance in this strain.

**Table 2.** CZA MIC of clinical isolates expressing ampD gene on a pUCP24 vector. Parent 2_C_ is the unevolved clinical isolate, Mutant 2_C_ is the 2_C_ isolate evolved in MEM.

| Strain | Parent (2<sub>C</sub>) | Mutant (2<sub>C</sub>) | Parent/<br>pUCP24 | Mutant/<br>pUCP24 | Parent/<br>pUCP*ampD* | Mutant/<br>pUCP*ampD* |
| --- | --- | --- | --- | --- | --- | --- |
| **CZA MIC,**<br>**mg/L** | 2 | 128 | 2 | 128 | 2 | 2 |

## Discussion

Treatment of *P. aeruginosa* infections is challenging due to the versatility of strains and their potential to develop antimicrobial resistance. In this multi-omics, multi-assay investigation of the *in vitro* evolution of resistance to meropenem (MEM) and ceftazidime-avibactam (CZA) in eight clinical *P. aeruginosa* isolates, we addressed these challenges both from the clinical and from the systems microbiology perspectives.

From the clinical perspective, our observations revealed that strains subjected to sub-inhibitory concentrations of MEM exhibited a considerably higher rate of and faster resistance evolution compared to those exposed to CZA. This suggests prolonged/inappropriate MEM use may accelerate resistance, potentially diminishing its therapeutic efficacy. In contrast, CZA demonstrated a lower propensity for resistance development.. These findings are in line with a recent study where *P. aeruginosa* isolates displayed greater increases of the mean MIC for MEM compared to CZA when subjected to increasing concentrations of the respective antibiotics (52).

Although cross-resistance to CZA appeared in fewer cases, continuous surveillance is important to inform and guide individualized treatment decisions for *P. aeruginosa* infections (6,7,9).

While six genes were found to be mutated independently by at least three evolved strains, these mutations were not clearly linked to resistance patterns (Figure 5). This reflects prior reports of overall lower mutation frequency in *P. aeruginosa*, underpinning that phenotypic variability may play a stronger role in the development of antibiotic resistance than adaptive mutations (53). Consistent with previous reports on high transcriptional heterogeneity in *P. aeruginosa* clinical isolates (53,54), we observed that the parental strain was the main driver of the transcriptional and proteomic profiles regardless of the antibiotic exposure.

Despite heterogeneous responses, machine learning approaches have previously been successful at predicting antibiotic resistance or virulence of clinical isolates based on genome or transcriptome features (34,55,56). Gene expression was more informative than mutation information for predicting resistance to meropenem and ceftazidime in *P. aeruginosa* (34). While our study’s small sample size limits building generalizable machine learning models, PLS-DA’s feature importance analysis helped us to identify candidate genes and proteins associated with resistance. Though potentially reflecting general drug stress response rather than direct resistance mechanisms, the identified genes and proteins could still serve as resistance and virulence biomarkers and inform combination treatments to suppress bacterial adaptation mechanisms and reduce antibiotic resistance.

The main categories of genes and proteins associated with antibiotic exposure include expected antibiotic resistance mechanisms such as membrane proteins, efflux pumps, and penicillin binding proteins, as well as a large group of metabolic genes involved in energy production and lipopolysaccharide metabolism. Energy metabolism was mostly downregulated, likely reflecting stress-induced reduced bacterial metabolism Conversely, lipopolysaccharide metabolism genes were generally upregulated, including those involved in alginate (PA0764 (*mucB*), PA0460) and lipid A biosynthesis (PA3552 (*arnB*), PA3554 (*arnA*)). Increased expression of arnA/B genes, along with mutations in their regulator *phoQ* are linked to resistance against polymyxin antibiotics and antimicrobial peptides (53–55), and overexpression of *arnA* and *arnB* were observed to be more common among *P. aeruginosa* isolates resistant to CZA, when compared to other anti-pseudomonal antibiotics (60). Similarly increased expression of alginate exopolysaccharide biosynthesis was previously observed in antibiotic resistant strains isolated from a cystic fibrosis patient (54), and also appeared predictive of ceftazidime or ciprofloxacin resistance in machine learning models (34,61). Since alginate production affects biofilm growth and decreases clearance by the host immune response (54), targeting alginate biosynthesis may be a strategy to complement antibiotic therapy.

The *mexAB-oprM* efflux system was preferentially upregulated in MEM-exposed strains. This system is regulated by the transcriptional repressors *mexR* and *nalD*, with additional indirect influence from *nalC* (62). Loss-of-function mutations in *mexR* and *nalD* are linked to increased resistance or elevated MICs to CZA (24,60). In our assays *nalD* mutations appeared in two MEM-exposed strains with significant upregulation of *mexA*, *mexB*, and *oprM*; however, these genes were also upregulated in three other strains with *nalD* downregulation, indicating *nalD* mutations are not the sole mechanism modulating the *mexAB-oprM* efflux system. Additionally, mutations in *oprD*, encoding the outer membrane porin *oprD*, resulted in substantial downregulation of this gene and were observed exclusively in MEM-exposed strains, further highlighting its role in carbapenem resistance (63).

In contrast, significant upregulation of *ampC* was primarily observed in strains exposed to CZA, suggesting that this antibiotic induces the cephalosporinase. Mutations in *dacB*, encoding penicillin-binding protein PBP4 and implicated in ampC derepression via ampR (30,59), were also found exclusively in CZA-selected strains.

A recent study detected AmpC overexpression in many of the isolates resistant to CZA (64). Mushtaq et al. (65) evaluated the role of *ampC* derepression in CZA resistance using 26 *Pseudomonas aeruginosa* strains with ampC mutations. Their findings showed that avibactam effectively reversed high-level ceftazidime resistance mediated by ampC, significantly reducing MICs in fully derepressed mutants. However, one strain with both high-level efflux activity and *ampC* derepression exhibited persistent resistance, highlighting the combined impact of efflux and *ampC* mutations on CZA resistance.

In our follow-up experiments with transposon insertion mutants in *P. aeruginosa* mPAO1 strain, only two out of fifteen mutants showed significant resistance changes, with CZA MIC decreasing by 25% or increasing up to three fold, though five other mutations were previously linked to altered resistance against various antibiotics (38) (Supplementary Table 16). Since transposon insertions may not replicate changes seen in evolved clinical isolates, this could explain the assay’s inability to reproduce resistance phenotypes.

To precisely address mutations found in genes associated with β-lactam resistance, we performed CRISPR/Cas9 genome editing of four of the mutated genes: *dacB*, *ampC*, *mexB* and *mexR*.

Mutations in *ampC* have been shown to enhance the ability of *ampC* to break down ceftazidime and to evade inhibition by avibactam more effectively (64). The introduction of a deletion within the *ampC,* identified in one of our strains that exhibited high resistance to CZA, resulted in only a slight increase in the CZA MIC level.

Previous studies identified a G115S mutation in *dacB* linked to *ampC* derepression (30), but its role in CZA resistance was unclear. While mutations slightly increased resistance overall, the most notable change was induced by a mutation in *dacB*, resulting in a four-fold increase in CZA MIC-levels, though allMICs remained below the clinically relevant threshold of MIC>8 mg/L. These results underline that antibiotic resistance mechanisms involve complex interactions beyond single-gene effects, emphasizing the need for a systems-level approach to understand their multifaceted nature.

Cloning experiments revealed that introducing a wild-type copy of the *ampD* gene had the most pronounced effect on restoring CZA sensitivity to mutant strains—highlighting its key role in acquiring CZA resistance. However, attempts to introduce mutations via CRISPR/Cas9 were unsuccessful despite several efforts; cloning served as an alternative approach but exaggerated *ampD*’s effect due to overexpression from high-copy vectors like pUCP24 (62) compared to chromosomally encoded *ampD*. Editing through CRISPR/Cas9, by contrast, would better reflect bacterial physiology because this approach enables modification of the endogenous *ampD* gene to be done in a precise manner that maintains native expression levels and regulatory control, thus more closely mimicking natural genetic variations in bacterial populations.

Together with previous reports, this study emphasizes the importance of comprehensive sampling and multi-omics analyses for identifying biomarkers associated with antibiotic resistance and inform combination therapies designed to suppress its evolution in clinicals isolates of *P. aeruginosa*.

## Materials and Methods

### Bacterial isolates and culture conditions

Eight clinical isolates of *Pseudomonas aeruginosa* were obtained from Kantonsspital St. Gallen hospital in Switzerland. The isolates were sourced from three different clinical sites, namely urine (n=4), sputum (n=3), and blood culture (n=1). For the multistep resistance selection process, bacterial cells were cultured in Luria Bertani (LB) Broth using Nunc™ CELLSTAR® 6-well plates (Greiner Bio-One GmbH, Frickenhausen, Germany). The cultures were maintained with shaking at 120 rpm on a microplate shaker (Edmund Bühler GmbH, Germany) at a temperature of 37°C.

Ceftazidime-avibactam (Zavicefta, Pfizer®) and meropenem (Meronem, Pfizer®) were procured from our hospital pharmacy at Kantonsspital St. Gallen. Antibiotic stock solutions were prepared by dissolving powder stocks in accordance with the manufacturer’s instructions. These solutions were then filter-sterilized through a 0.2 µm filter and stored at -20°C until used.

### MIC determination / antimicrobial susceptibility testing

Minimal inhibitory concentrations (MICs) for several antibiotics were determined by broth microdilution in Mueller–Hinton (MH) broth using the Sensititre GNX2F plates (Thermo Fisher scientific, East Grinstead, United Kingdom). MICs for meropenem (MEM) and ceftazidime-avibactam (CZA) were further determined by Etest-strips (bioMérieux, Chemin de l’Orme, France) and broth microdilution method (Supplementary Table 4).

MICs for ceftolozane-tazobactam (TOL-TAZ) were determined by broth microdilution in Mueller Hinton (MH) broth using ComASP™ Ceftolozane-tazobactam (Liofilchem, Brunschwig, Basel, Switzerland) and by Etest (bioMérieux, Chemin de l’Orme, France). The MIC determination adhered to the latest European Committee on Antimicrobial Susceptibility Testing (EUCAST) recommendations (66), with values interpreted in accordance with the 2022 EUCAST criteria (version 12.0).

### Evolution for antimicrobial resistance (multi-step resistance selection)

Eight representative strains were selected from each of the original eight *Pseudomonas aeruginosa* isolates. This process involved isolating a single colony forming unit (CFU), which was subsequently streaked onto a fresh Mueller Hinton (MH) agar plate. From each of these plates, three distinct colonies were carefully chosen to serve as parental strains for subsequent passage experiments. This resulted in 24 strains being passaged in each of the antibiotics (24 strains in CZA, 24 strains in MEM). For each individual strain, separate minimal inhibitory concentration (MIC) measurements were conducted for ceftazidime-avibactam (CZA) and meropenem (MEM). To initiate the experiments, a single colony of each strain was selected after overnight growth at 37°C on an MH agar plate. Subsequently, each chosen colony was grown in 5 ml of Luria Bertani (LB) broth at 37°C for 18 hours without the presence of antibiotics (overnight culture). The strains were then concurrently exposed to sub-inhibitory concentrations, equivalent to one-fourth of the MIC values for each respective strain, of either CZA (n=3) or MEM (n=3). In brief, 5 ml of LB broth containing the corresponding antibiotic at a concentration of one-fourth of the MIC was inoculated with 25 µl of the overnight broth. The cultures were then agitated on a shaker under the previously described conditions, maintained at 37°C for a duration of 24 hours. Subsequently, a daily transfer of 25 µL from the overnight culture was made to new LB broth containing CZA or MEM concentrations corresponding to one-fourth of the measured MIC. Following six consecutive days under the consistent influence of the same antibiotic concentration, the minimal inhibitory concentrations (MICs) were reassessed using the Etest. Based on the outcomes of these re-evaluated MICs, modifications were implemented to the antibiotic concentration for the subsequent cycle of passage.

Aliquots of 50 µl of the bacterial cultures were sampled after each six passages and kept in 1 ml 10% Glycerol at −80°C for further molecular investigations. Furthermore, a subset of strains was chosen at random and underwent Matrix-Assisted Laser Desorption/Ionization Time-of-Flight (MALDI-TOF) analysis to ensure the absence of contaminants. To thoroughly assess the trajectory of resistance development and cross-resistance, the aforementioned protocol was replicated over a total of 18 sequential days. To comprehensively investigate the molecular mechanisms underlying resistance, those specific strains that had not exceeded the clinical breakpoint defined for each respective antibiotic, as specified by the EUCAST guidelines (67), underwent additional exposure to sub-inhibitory concentrations of the corresponding antibiotics. This exposure was continued until the strains developed resistance or for a maximum of 42 days. Concurrently, an identical procedure was conducted in the absence of antibiotic pressure, serving as negative controls for comparison.

### Whole Genome sequencing

DNA extractions were conducted following the protocols outlined in the handbooks of two distinct kits: the QIAamp DNA Mini Kit (QIAGEN, Hilden, Germany) was employed for Illumina sequencing, while the QIAGEN Genomic-tip 100/G + Genomic DNA Buffer Set (QIAGEN, Hilden, Germany) kits were used for the parental strains that underwent PacBio sequencing. Whole genome sequencing was carried out by Novogene (Cambridge, United Kingdom) and the Genomics Facility of the Department of Biosystems Science and Engineering of ETH Zürich (Basel, Switzerland).

For PacBio sequencing, a 10 kb SMRTbell library was prepared using the SMRTbell Template Prep Kit 1.0-SPv3 (Pacific Biosciences of California, USA). The qualified high-molecular-weight DNA was fragmented to approximately 10 kb, followed by processes such as damage repair, end repair, and adapter ligation. Subsequently, size selection was accomplished using a Size-Selection System.

The SMRTbell-Polymerase Complex was prepared utilizing the Sequel™ Binding Kit 2.0 (Pacific Biosciences of California, USA) and subsequently subjected to sequencing on a Sequel SMRT Cell (Pacific Biosciences of California, USA).

For Illumina sequencing, DNA samples were processed for library preparation following the manufacturer’s recommendations of the NEBNext® DNA Library Prep Kit (New England BioLabs, US). Index codes were incorporated for each sample. In summary, genomic DNA was randomly fragmented to a size of 350 bp. The DNA fragments underwent end polishing, A-tailing, ligation with adapters, size selection, and further PCR enrichment. Subsequently, PCR products were purified using the AMPure XP system, their size distribution was assessed using the Agilent 2100 Bioanalyzer (Agilent Technologies, CA, USA), and quantification was performed using real-time PCR. The library was then sequenced on a NovaSeq 6000 S4 flow cell employing the PE150 strategy.

### Mutation identification

Fasta files from the whole genome sequencing were analyzed with the snippy perl package (version 4.6.0) (https://github.com/tseemann/snippy) to call the single nucleotide polymorphisms (SNPs) and other types of mutations using each parent strain genome sequenced before treatment as a reference. Other than options specifying the input files and output folder, snippy was called with default parameters and force and report flags and parallelised on 16 cores.

### Sample preparation for transcriptomic profiling

The gene expression profiles of the resistant strains and the corresponding susceptible parent strains were analyzed with RNA sequencing involving three technical replicates for each condition. For RNA extraction, the RNAprotect Bacteria Reagent & RNeasy Mini kits (QIAGEN, Hilden, Germany) were utilized, following the RNAprotect® Bacteria Reagent Handbook protocols 4 (Enzymatic Lysis and Proteinase K Digestion of Bacteria) and 7 (Purification of Total RNA from Bacterial Lysate Using the RNeasy Mini Kit). An optional on-column DNase digestion was performed using the RNase-Free DNase set (QIAGEN, Hilden, Germany), and ribosomal RNA was depleted with the NEBNext® rRNA Depletion Kit (New England BioLabs, Allschwil, Switzerland). The resulting RNA was fragmented, reverse transcribed, and prepared for sequencing on an Illumina NextSeq 500 platform. Sequencing comprised 150 bp single-end reads with a target of 5-10E6 reads per strain.

### Protein extraction

3ml of OD 1 bacteria culture was pelleted by centrifugation (14’000 x g for 5 minutes) and was washed 4x with 1x PBS and lysed with 100 µl of buffer (4% (w/v) SDS, 100 mM Tris/HCL pH 8.2, 0.1M DTT + phosphoSTOP (Roche Diagnostics GmbH, Mannheim, Germany). To each sample 100 µl of lysis buffer (4% Sodium dodecyl sulfate (SDS) in 50 mM triethylammonium bicarbonate buffer (TEAB) were added.

The samples were frozen in liquid nitrogen for 1 min and subsequently sonicated (sonication bath) for 2 min. These freeze/thaw cycles were repeated for a total of three times. To each sample 12.5 Units of benzonase (Sigma-Aldrich) were added and incubated for 15 min at RT, 700 rpm in a thermoshaker (Eppendorf).

The samples were treated with High Intensity Focused Ultrasound (HIFU) for 1 minute at an ultrasonic amplitude of 85% followed by further extraction using a tissue homogenizer (TissueLyser II, QIAGEN) for 4 min at 30 Hz. After another 1 min of HIFU the samples were centrifuged at 20’000 x g for 10 min. The protein concentration was determined using a Lunatic (Unchained Labs) instrument. For each sample 50 µg of protein were taken and reduced with 2 mM TCEP (tris(2-carboxyethyl)phosphine) and alkylated with 15 mM iodoacetamide at 60°C for 30 min.

### Protein digestion

The sp3 protein purification, digest and peptide clean-up was performed using a KingFisher Flex System (Thermo Fisher Scientific) and Carboxylate-Modified Magnetic Particles (GE Life Sciences) (68,69). Beads were conditioned following the manufacturer’s instructions, consisting of 3 washes with water at a concentration of 1 ug/ul. Samples were diluted with an equal volume of 100% ethanol (50% ethanol final concentration). The beads, wash solutions and samples were loaded into 96 deep well- or micro-plates and transferred to the KingFisher.

Following steps were carried out on the robot: collection of beads from the last wash, protein binding to beads (14 min), washing of beads in wash solutions 1-3 (80% ethanol, 3 min each), protein digestion (offline from the KingFisher, overnight at RT with a trypsin:protein ratio of 1:50 in 50 mM TEAB) and peptide elution from the magnetic beads (water, 6 min). The digest solution and water elution were combined and dried to completeness. Afterwards the peptides were re-solubilized with 40 µl of 3% acetonitrile, 0.1% formic acid for MS analysis.

### Proteomic analysis with liquid chromatography-mass spectrometry

Mass spectrometry (MS) analysis was performed on an Orbitrap Fusion Lumos (Thermo Scientific) equipped with a Digital PicoView source (New Objective) and coupled to a M-Class UPLC (Waters). Solvent composition of the two channels was 0.1% formic acid for channel A and 0.1% formic acid, 99.9% acetonitrile for channel B. For each sample 2 μl of peptides were loaded on a commercial MZ Symmetry C18 Trap Column (100Å, 5 µm, 180 µm x 20 mm, Waters) followed by nanoEase MZ C18 HSS T3 Column (100Å, 1.8 µm, 75 µm x 250 mm, Waters). The peptides were eluted at a flow rate of 300 nl/min. After an initial hold at 5% B for 3 min, a gradient from 5 to 24% B in 80 min and 36% B in 10 min was applied. The column was washed with 95% B for 10 min and afterwards the column was re-equilibrated to starting conditions for additional 10 min. Samples were acquired in a randomized order. The mass spectrometer was operated in data-dependent mode (DDA) acquiring a full-scan MS spectrum (300−1’500 m/z) at a resolution of 120’000 at 200 m/z after accumulation to a target value of 500’000. Data-dependent MS/MS were recorded in the linear ion trap using quadrupole isolation with a window of 0.8 Da and HCD fragmentation with 35% fragmentation energy. The ion trap was operated in rapid scan mode with a target value of 10’000 and a maximum injection time of 50 ms. Only precursors with intensity above 5’000 were selected for MS/MS and the maximum cycle time was set to 3 s. Charge state screening was enabled. Singly, unassigned, and charge states higher than seven were rejected. Precursor masses previously selected for MS/MS measurement were excluded from further selection for 20 s, and the exclusion window was set at 10 ppm. The samples were acquired using internal lock mass calibration on m/z 371.1012 and 445.1200. The mass spectrometry proteomics data were handled using the local laboratory information management system (LIMS) (70) and all relevant data have been deposited to the ProteomeXchange Consortium via the PRIDE (http://www.ebi.ac.uk/pride) partner repository with the data set identifier PXD044998.

### Protein identification and label free quantification

The acquired raw MS data were processed by MaxQuant (version 1.6.2.3), followed by protein identification using the integrated Andromeda search engine (71). Spectra were searched against a *Pseudomonas aeruginosa* reference proteome (version from 2021-01-12), concatenated to its decoyed fasta database and common protein contaminants. Carbamidomethylation of cysteine was set as fixed modification, while methionine oxidation and N-terminal protein acetylation were set as variable. Enzyme specificity was set to trypsin/P allowing a minimal peptide length of 7 amino acids and a maximum of two missed-cleavages. MaxQuant Orbitrap default search settings were used. The maximum false discovery rate (FDR) was set to 0.01 for peptides and 0.05 for proteins. Label free quantification was enabled and a 2 minutes window for match between runs was applied. In the MaxQuant experimental design template, each file is kept separate in the experimental design to obtain individual quantitative values. Protein fold changes were computed based on Intensity values reported in the proteinGroups.txt file. The proteinGroups.txt file generated by MaxQuant was used for downstream analysis to compute a moderated t-test (72) for all proteins quantified with at least 2 peptides employing the R package limma (73). Protein abundances were normalised using quantile normalisation. A set of functions implemented in the R package SRMService (74) was used to filter for proteins with 2 or more peptides allowing for a maximum of 4 missing values, and to normalize the data with a modified robust z-score transformation and to compute p-values using the t-test with pooled variance. If all measurements of a protein are missing in one of the conditions, a pseudo fold change was computed replacing the missing group average by the mean of 10% smallest protein intensities in that condition.

### Transcriptomics data analysis

Quantification of transcriptomic data was performed by using Trimmomatic (version 0.39) (75) followed by Salmon (version 1.10.1) (76) packages. DESeq2 (version 1.38.0) (77) R package was used to perform normalisation and differential expression analysis, first by importing the data generated by Salmon package using tximport R package (version 1.26.0) (78), then by running DESeqDataSetFromTximport and DESeq functions. The aforementioned tools were run on all samples using a Snakemake pipeline (58). Differential analysis was performed for each strain comparing expression levels in each evolved strain (on CZA and MEM) to the parent strain expression, as well as between CZA- and MEM-evolved strains of the same parent. Furthermore, to filter the results from PLS-DA feature importance analysis, all the strains were grouped based on their MEM- or CZA-resistance levels with the threshold of MIC=32 mg/L indicating high resistance. Differential analysis was performed between all sensitive versus resistant strains for each antibiotic (for MEM: all parent strains and CZA-exposed 2_C_, 3_C_, 4_C_, 5_B_ and 6_C_ strains versus CZA-exposed 1_B_ and all MEM-exposed strains; for CZA: all parent strains, CZA-exposed 3_C_ and 6_C_ and MEM-exposed 2_C_, 3_C_, 4_C_ and 5_B_ versus CZA-exposed 1**_B_**, 2_C_, 4_C_, 5_B_ and MEM-exposed 1_B_ and 6_C_ strains), as well as all sensitive versus resistant exposed to the same antibiotic (for MEM: CZA-exposed 2_C_, 3_C_, 4_C_, 5_B_ and 6_C_ strains versus CZA-exposed 1_B_ strain; for CZA: CZA-exposed 3_C_ and 6_C_ versus CZA-exposed 1_B_, 2_C_, 4_C_, 5_B_ strains, or MEM-exposed 2_C_, 3_C_, 4_C_ and 5_B_ strains versus MEM-exposed 1_B_ and 6_C_ strains). DESeq2 utilises Wald test to compare expression levels between conditions with Benjamini-Hochberg correction for multiple hypothesis testing. All the processing of the data was executed on the EMBL Heidelberg HPC cluster (80).

### Partial least squares discriminant analysis

PLS-DA was performed by using the function PLSRegression from scikit-learn python package (version 1.3.0) to predict whether CZA or MEM resistance was high (1) or low (0). Threshold of MIC=32 mg/L was used to define high resistance. First set of models used all of the transcriptomic and proteomic data for fitting (n=51 samples), to determine the separation of samples with high and low MIC values in the lower dimensional space. The second set of models used a leave-one-strain-out strategy - data related to one of the strains left out of the training set (n=6-9 samples), then this data was used as a test set to determine the accuracy of the predictions for the model trained on all the other strains (n=42-45 samples). To determine a set of the most important features (genes or proteins) contributing to the separation of the samples with high and low MIC values, top one hundred features by absolute values of weights were selected from each leave-one-strain-out model for the first and the second PLS factor (200 features per model in total).

### Gene ontology enrichment analysis

Gene ontology terms were determined by querying PantherDB REST API (https://pantherdb.org/services/oai/pantherdb/enrich/overrep) with a set of genes or proteins that resulted from differential expression and differential abundance analyses and PLS-DA analysis. Significantly up- or down-regulated (FDR < 0.05 and abs(log2FC) > 1) genes and proteins from differential expression and differential abundance analyses were passed to the PantherDB to perform over-representation analysis. Similarly, genes and proteins determined by PLS-DA as contributing to the top two hundred weights of leave-one-strain-out models and appearing in at least five models were passed to PantherDB API. Over-representation analysis done by PantherDB service assumes that all *P. aeruginosa* genes are present in the reference list.

### Sequence typing and phylogenetic analysis

To have the genetic background of the strains in this study compared with each other and with genomes of other publicly available strains, sequence typing analysis and phylogenetic analysis were performed. Sequence typing was performed using PubMLST web service (https://pubmlst.org) (81). Phylogenetic analysis was performed using Mashtree (version 1.4.6) (78), first comparing just the strains in this study, then comparing them to strains with complete genomes available from a database of pseudomonas genomes (https://pseudomonas.com) (82–84) and matplotlib (version 3.8.3) (85).

### Comparison to public datasets

Several publicly available datasets were used for comparison with the results of the differential expression analysis conducted in this study. Dong *et al.* (*2024*) dataset (86) was processed from raw reads with the same pipeline as for the dataset in this study, specifically by using Salmon (version 1.10.1) (76) followed by DESeq2 (version 1.38.0) (77). Khaledi *et al.* (2020) dataset, consisting of a counts table, was processed with DESeq2 as well, by running DESeqDataSetFromMatrix and then DESeq functions. Additionally, two datasets, Alam 2023 (87) and Castanheira 2022 (35), had already undergone differential expression analysis. Significantly up- or down-regulated genes from each dataset were compared with those from this study on a per-strain basis. This comparison was performed by calculating the ratio of the size of the intersection of two given sets to the size of each set and applying Fisher’s exact test to the intersection. P-values were subsequently adjusted with the Benjamini-Hochberg procedure.

### Data selection and visualisation for the summary plot

A data-driven approach was employed to select the genes and proteins shown in Figure 5 from mutation analysis, differential analysis and PLS-DA. From mutation analysis, genes with mutations present in at least two strains were selected. From differential analysis, genes and proteins significantly up- or downregulated (abs(log2(fold change))>=1 in either of the omics, FDR<0.05 in both omics) by at least five strains from transcriptomics, four strains from proteomics and three strains from both were selected. From PLS-DA, genes and proteins that appear in the top two hundred features in at least five leave-one-strain-out models and additionally significantly differentially abundant between the groups of sensitive and resistant strains (see section *Transcriptomics data analysis* for details) were selected. The grouping of the final list of genes and proteins, obtained by combining these sets of genes and proteins, was performed by ChatGPT by providing the list of locus tags and using the prompt: “can you group the following list of genes based on their functions” followed by manual checks and corrections.

### MIC measurement of the selected mutant strains of P. aeruginosa mPAO1

*P. aeruginosa* mPAO1 parent strain and its transposon insertion mutants were obtained from the *P. aeruginosa* Two-Allele Library of the Manoil Lab (88). Bacteria were activated on Plate-Count-Agar (Sigma Aldrich, Steinheim, Germany). Single colonies from the plates were picked up and inoculated in 25 mL Müller Hinton Broth (MHB, Sigma Aldrich, Steinheim, Germany) and cultivated overnight at 37°C and 160 rpm. 250 μl of the overnight culture was added to 25 mL MHB and further incubated for 3 hours to obtain the exponentially growing cells. The final culture was diluted to an optical density (OD) of 0.1 measured at 600 nm, which corresponds to approximately 10^8^ CFU/ml, of which 500 µl was transferred to 50 ml MHB and mixed well by vortexing. Antibiotic stock solutions for MEM were prepared as mentioned above. CZA for micro broth dilution was prepared with ceftazidime (Sigma Aldrich, Steinheim, Germany) and avibactam (Sigma Aldrich, Steinheim, Germany), the latter with a fixed concentration of 4 mg/l per well.

The MIC assays of the MEM and CZA were performed in 96 well plates (BRAND® microplate BRANDplates®, Sigma Aldrich, Steinheim, Germany) covered with an air-permeable foil (Breathe-Easy sealing membrane Sigma Aldrich, Steinheim, Germany) and incubated at 37°C without shaking. All assays were done in MBH medium supplemented with increasing concentrations of antibiotics, and inoculated with mPAO1 mutants of 5*10^5^ CFU/ml. For each condition, four replicates were included. After 24 h at 37°C, the bacterial growth was measured at OD 600 nm.

### Cloning of ampD gene in the evolved CZA-resistant strain

The wildtype *ampD* allele was amplified from the parental 2_C_ isolate using primers ampD-Fw (5′-GATGATGAGCTCGCGCCGCTGGATTAGAGT-3’) and ampD-Rev (5′-

GATGATGGATCCCAGCAGCAACACCAGGAAC-3′), and cloned into vector pUCP24, before transformation into *E. coli* Top10. Transformants were selected on plates supplemented with gentamicin (25 mg/L). Successful transformants were confirmed by PCR amplification and sequencing of the alleles, then transformed into the parent and the CZA-evolved mutant 2_C_ strains. Susceptibility testing was performed by disc diffusion, and MICs were determined by broth microdilution according to EUCAST guidelines.

### Crispr/Cas9 genome editing

CRISPR/Cas9 genome editing was performed using the two plasmid (pCasPA/pACRISPR) method developed by Chen et al. (89) using the λ-Red recombination system.

sgRNA oligos were designed with CRISPOR (90) and the overhangs added according to Supplementary Table 16.

Oligos were phosphorylated and annealed by mixing 35 µl H2O, 5 µl 10x T4 Polynucleotide Kinase Buffer (New England BioLabs, Ipswich, MA, USA), 5 µl fresh 10mM ATP, 1 µl T4 polynucleotide kinase (New England BioLabs, Ipswich, MA, USA), 2 µl of each oligo (50 µM). After one hour incubation at 37°C, 2.5 µl 1M NaCl were added, the mix heated to 95°C for 3 minutes and then cooled to 25°C by decreasing the temperature -0.1°C per second. Annealed oligos were diluted 1:20 and 1 µl mixed with 1.5 µl H2O, 5 µl 2x Quick Ligase Buffer (NEB), 20nM pACRISPR plasmid, 0.5µl Quick Ligase (400U/ µl), 0.5 µl BsaI-HFv2 (NEB) and 0.5 µl fresh 10mM ATP. Reaction was kept in a thermocycler for 25 cycles (37°C for 3 min, 16°C for 4 min), then 80°C for 15 min, and finally cooled to 10°C. The product was transformed into zymo mix and go (Takara, Mountain View, CA, USA) competent *E. coli* and selected on 50 mg/l carbenicillin plates. Insertion of the sgRNA sequence was verified by sequencing (sgSeq-F primer). Primer for the repair 500 bp arms were designed with the Takara In-Fusion Cloning Primer Design Tool to contain the desired restriction sites and containing the desired nucleotide change and a synonymous change of the PAM sequence. Repair arms were inserted in between the XhoI and XbaI restriction site of the pACRISPR plasmid following the Takara In-Fusion PCR Cloning systems protocol and insertion verified by PCR and sequencing. *P. aeruginosa* mPAO1 was made electrocompetent by diluting an overnight culture 1:100 in fresh LBB and incubating them at 37°C until OD600 of ∼0.5. Cells were chilled on ice for 30 minutes, then were harvested by centrifugation at 3300 rpm for 5 min and pellet washed twice with 20 ml of ice-cold 10% glycerol and then resuspended in 1 ml 10% glycerol. 50 µl electrocompetent cells were mixed with up to 5 µl of pCas plasmid and electroporated in a 1 mm cuvette (1.8kV, 200 Ω, 25 µF). Immediately after the pulse 1 ml LBB was added and the mix was incubated for 90 minutes at 37° C, 200 rpm followed by selection on 30 mg/l tetracycline plates.

Afterwards one colony harbouring the pCasPA plasmid was prepared as electrocompetent cells, same as above except for the addition of 2 mg/L total concentration L-arabinose at OD 0.3. pACRISPR with the sgRNA sequence and repair arms were mixed and electroporation followed with the same conditions. 30 mg/l tetracycline and 100 mg/l carbenicillin were used for selection.

Transformants were plasmid cured by growing them in the absence of antibiotics until evident growth and then plating a 1:10^4^ dilution on 5% sucrose LBA. The PCR amplified region of the CRISPR site of Plasmid cured bacteria was sequenced to confirm the CRISPR event.

## Data availability

The genomics and transcriptomics data have been deposited in the European Nucleotide Archive (ENA) repository https://www.ebi.ac.uk/ena/browser/view/PRJEB76120. Proteomics data have been deposited to the ProteomeXchange Consortium via the PRIDE (http://www.ebi.ac.uk/pride) repository with the data set identifier PXD044998. Supplementary tables and supporting data are available in the following Zenodo archive https://zenodo.org/uploads/15235346.

## Code availability

Code to reproduce all the analysis in this work can be found in the following git repository https://git.embl.de/grp-zimmermann-kogadeeva/PseudomonasAntibioticsResistance.

## Acknowledgements

We thank the members of the Zimmermann-Kogadeeva lab for helpful discussions; the Functional Genomics Center Zurich for technical assistance with proteomics measurements, Stefanie Altenried and Flavia Zuber at Empa for their assistance with analysing the mPAO1 insertion mutants, Valentin Gisler from Cantonal Hospital Aarau for providing clinical isolates, the Genomics Facility Basel (Department of Biosystems Science and Engineering, ETH Zürich) for WGS and RNAseq analysis. We also thank the EMBL IT Services staff for managing and provision of access to the HPC resources.

This work has been presented at the 34th Congress of the European Society of Clinical Microbiology and Infectious Diseases in Barcelona, Spain (EO243) and the Joint Annual Meeting of the Swiss Society of Infectious Diseases and Microbiology in Bern, Switzerland (O077).

## Author Contributions

Study and experiment design: BBF and MZ-K. Data analysis: BJB, SS, MZ-K, AB. *In vitro* experiments: AB, JF, QR, NRI. Clinical isolate work: AB, BBF, AE. Manuscript writing: BJB, MZ-K, BBF, AB.

Manuscript editing: BJB, AB, SS, NRI, QR, JF, AE, MZ-K, BBF. Funding acquisition: BBF, MZ-K. All authors read and approved the manuscript.

## Statement on using AI tools

In the preparation of this research paper, we utilised generative AI tools (ChatGPT 4o and perplexity.ai) to enhance the clarity and coherence of our writing, including this section. We carefully reviewed and edited the AI-generated recommendations to maintain the integrity and authenticity of our work. Generative AI-tools were not used to produce new content, but only to rephrase human-written text to improve its clarity. ChatGPT 4o was also used to suggest the clustering of genes based on their functions in Figure 5 (described in the Methods section), which was subsequently edited by the authors.

---

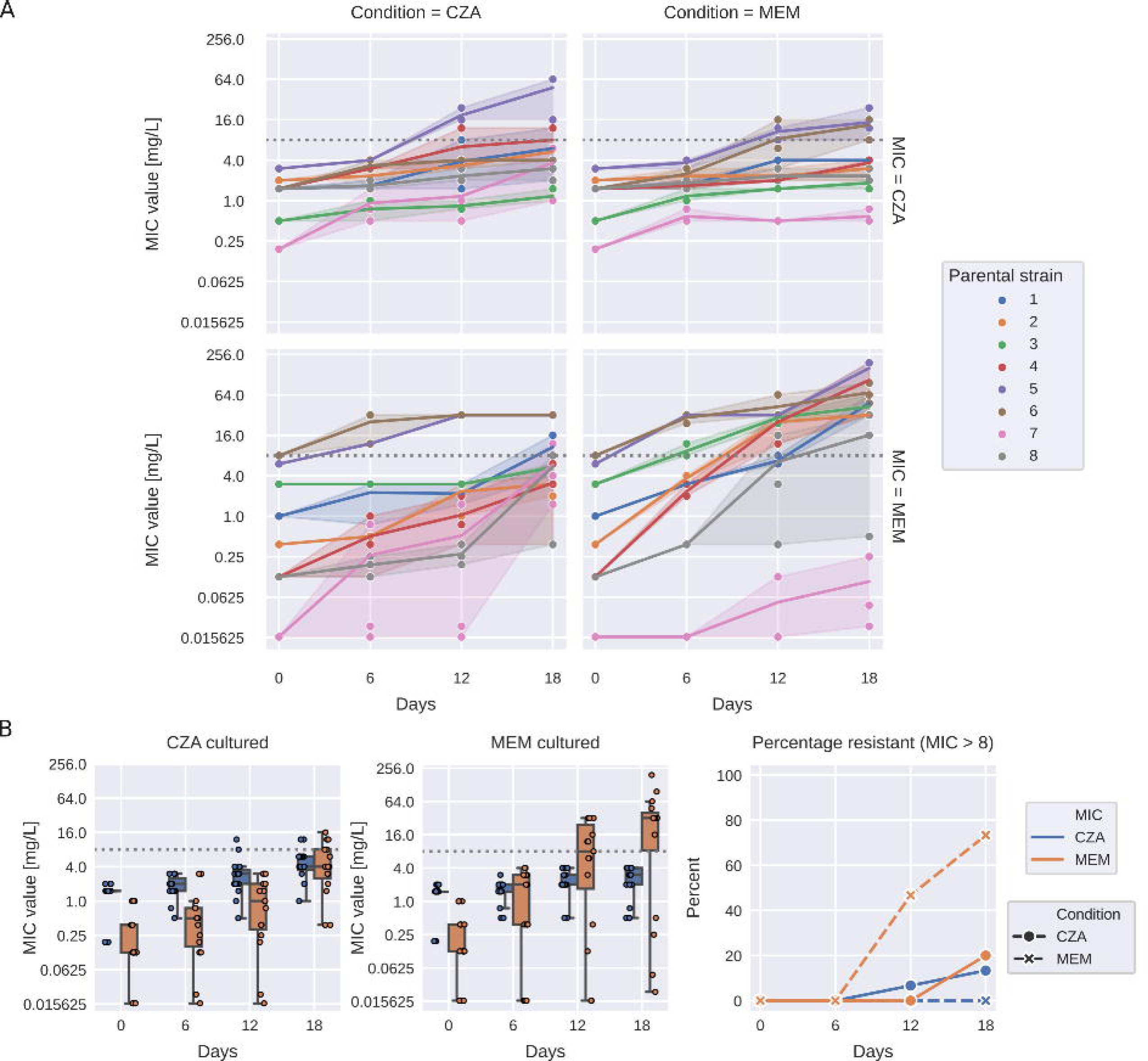

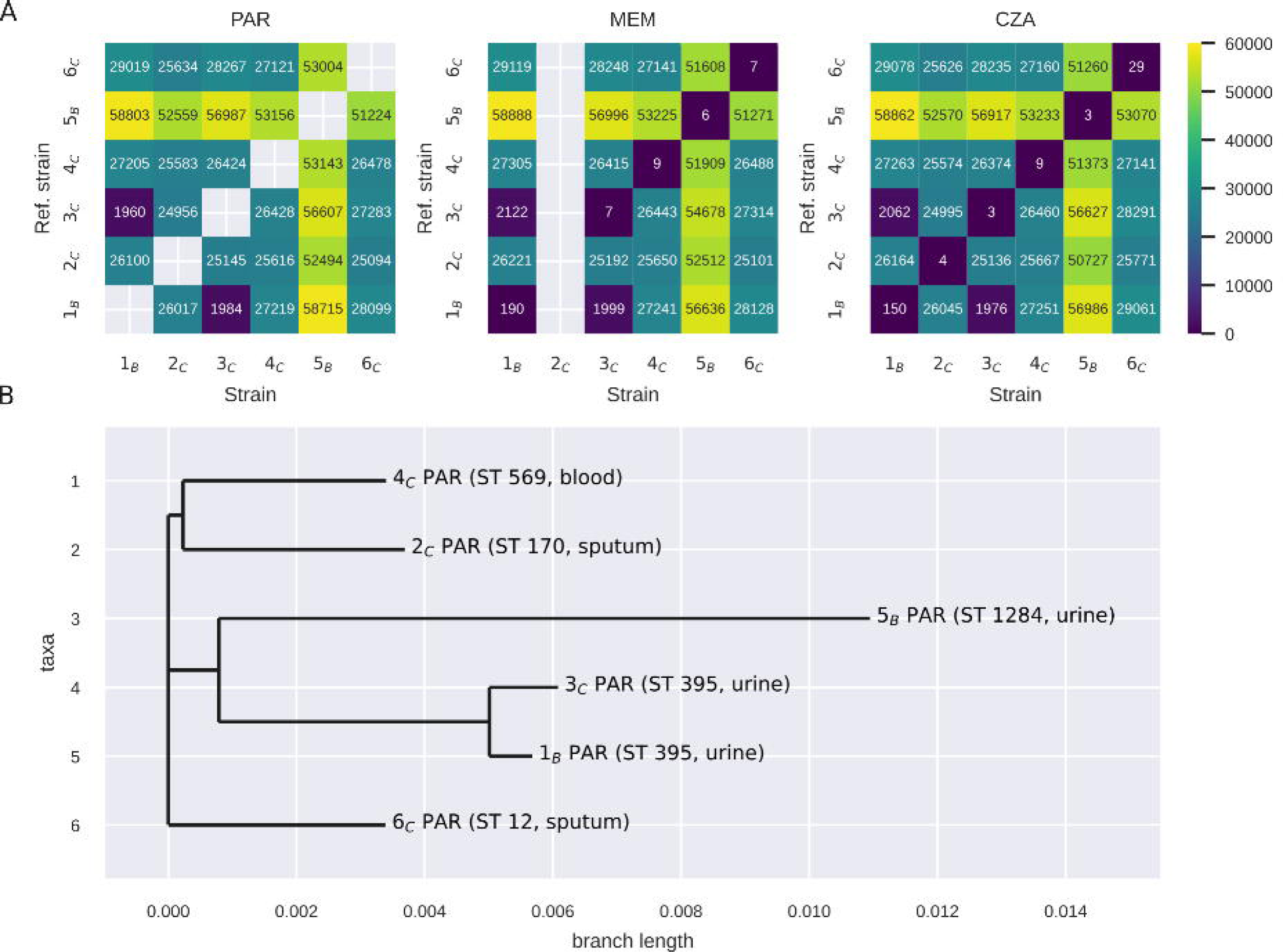

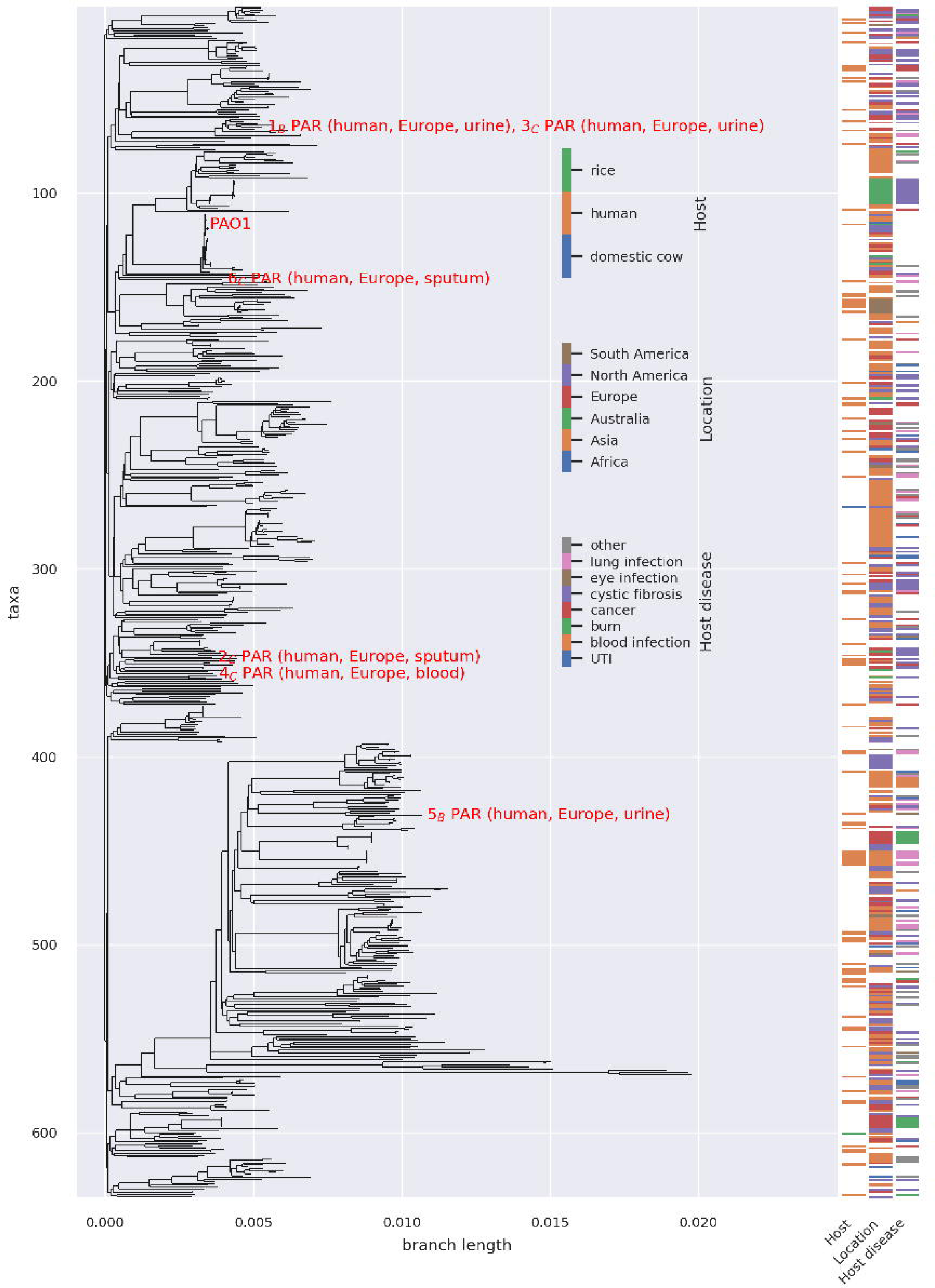

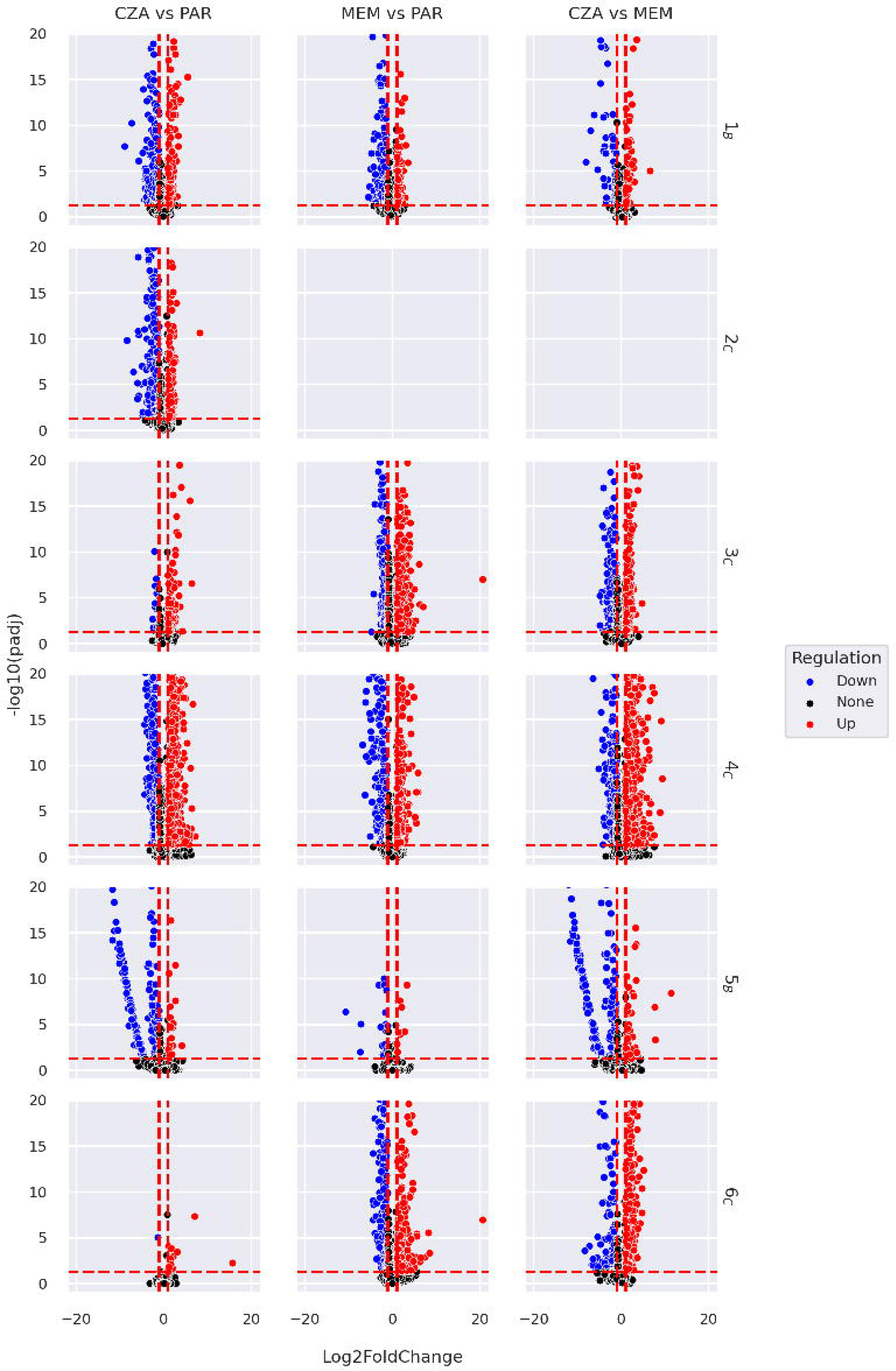

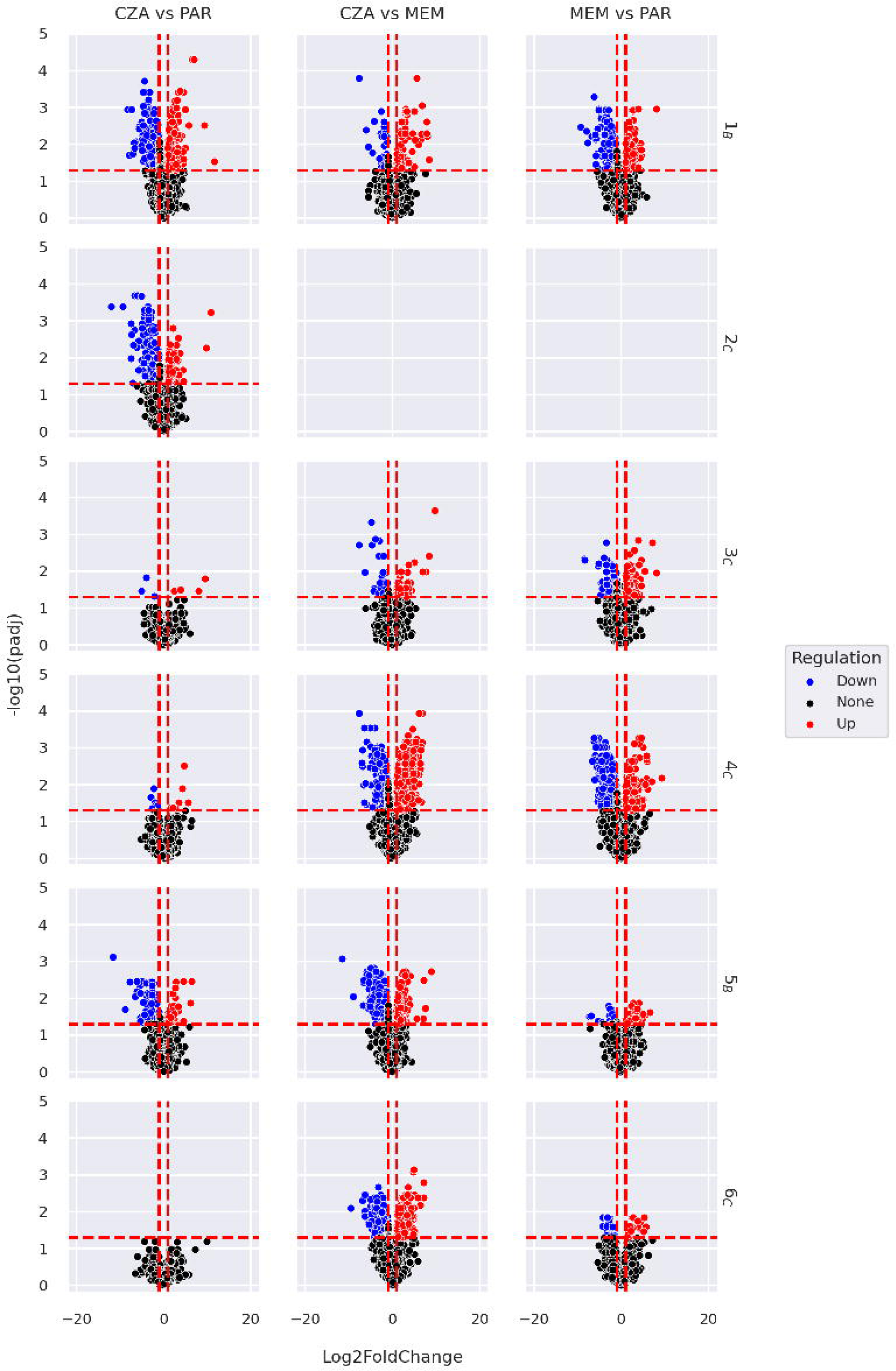

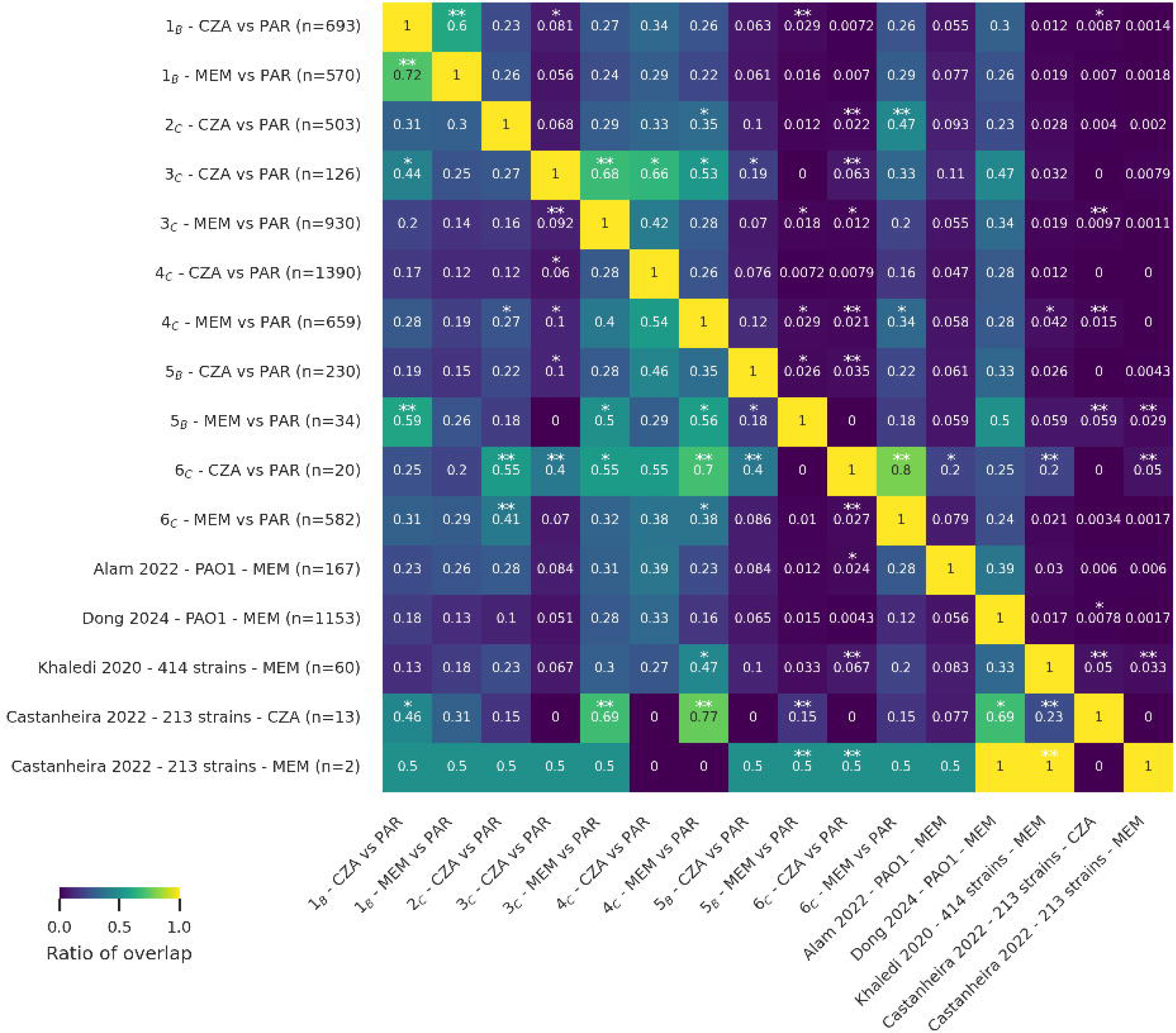

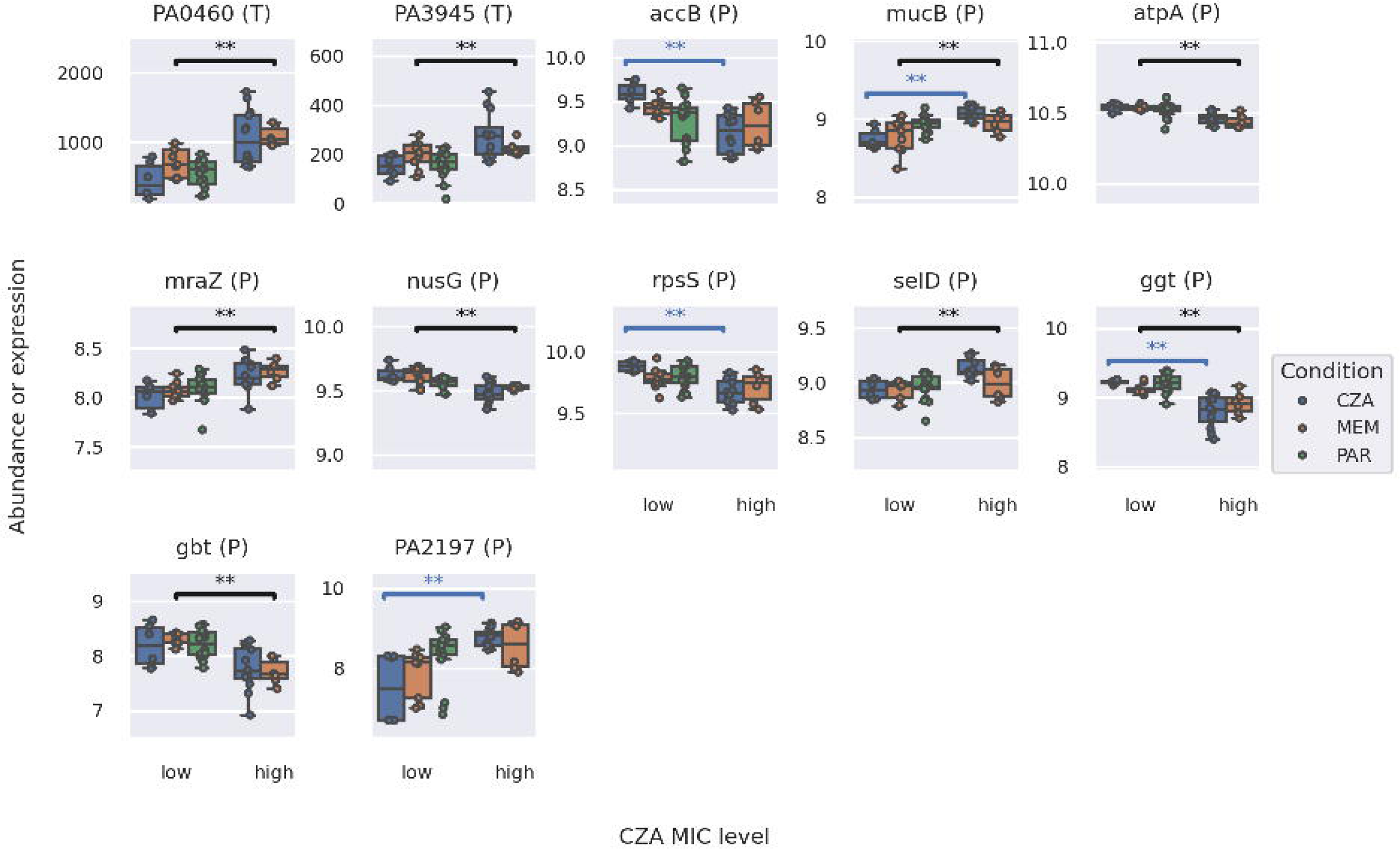

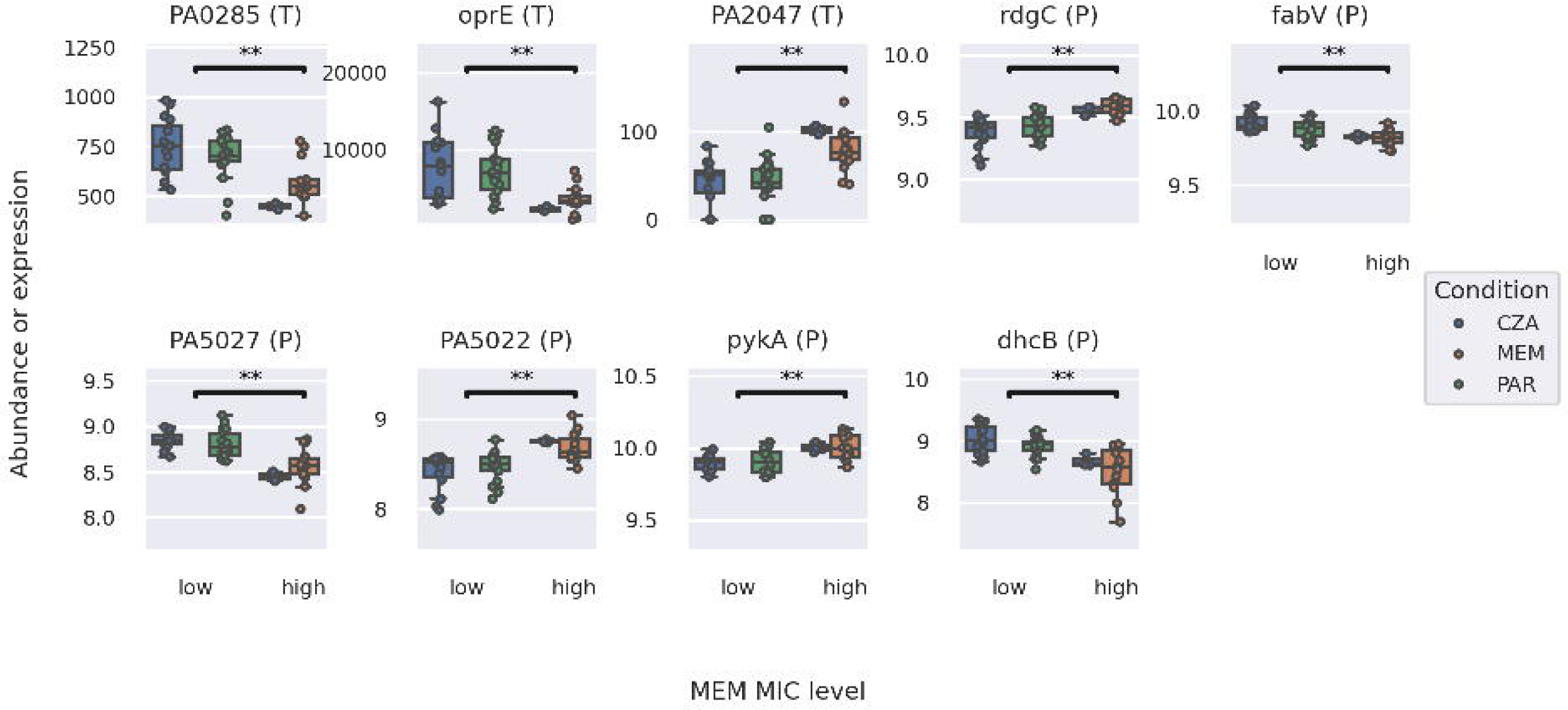

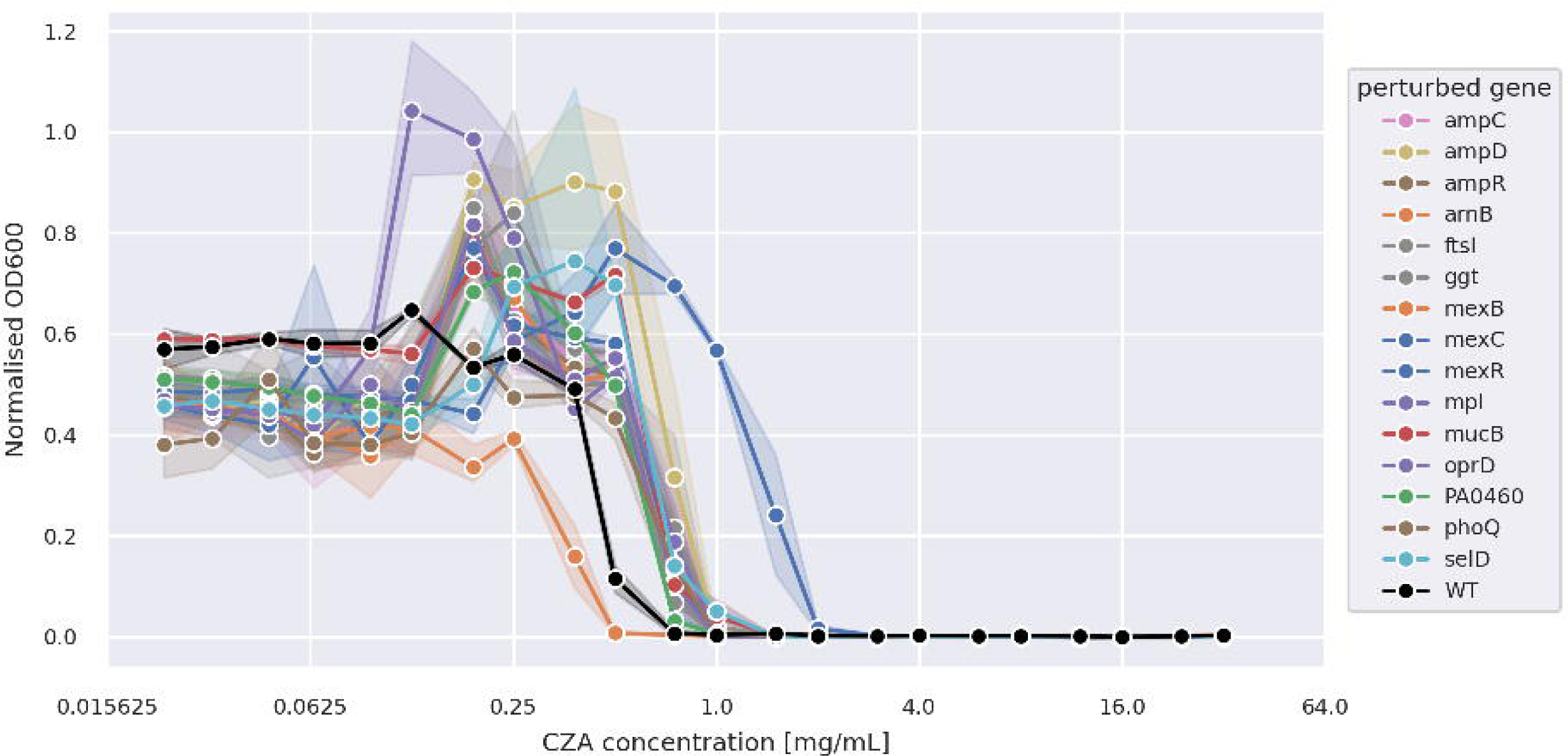

