## Supplementary figures and table legends and supplementary table 16 for "Multi-omics profiling of cross-resistance between ceftazidime-avibactam and meropenem identifies common and strain-specific mechanisms in *Pseudomonas aeruginosa* clinical isolates"

**Supplementary information**

**Supplementary Figures**

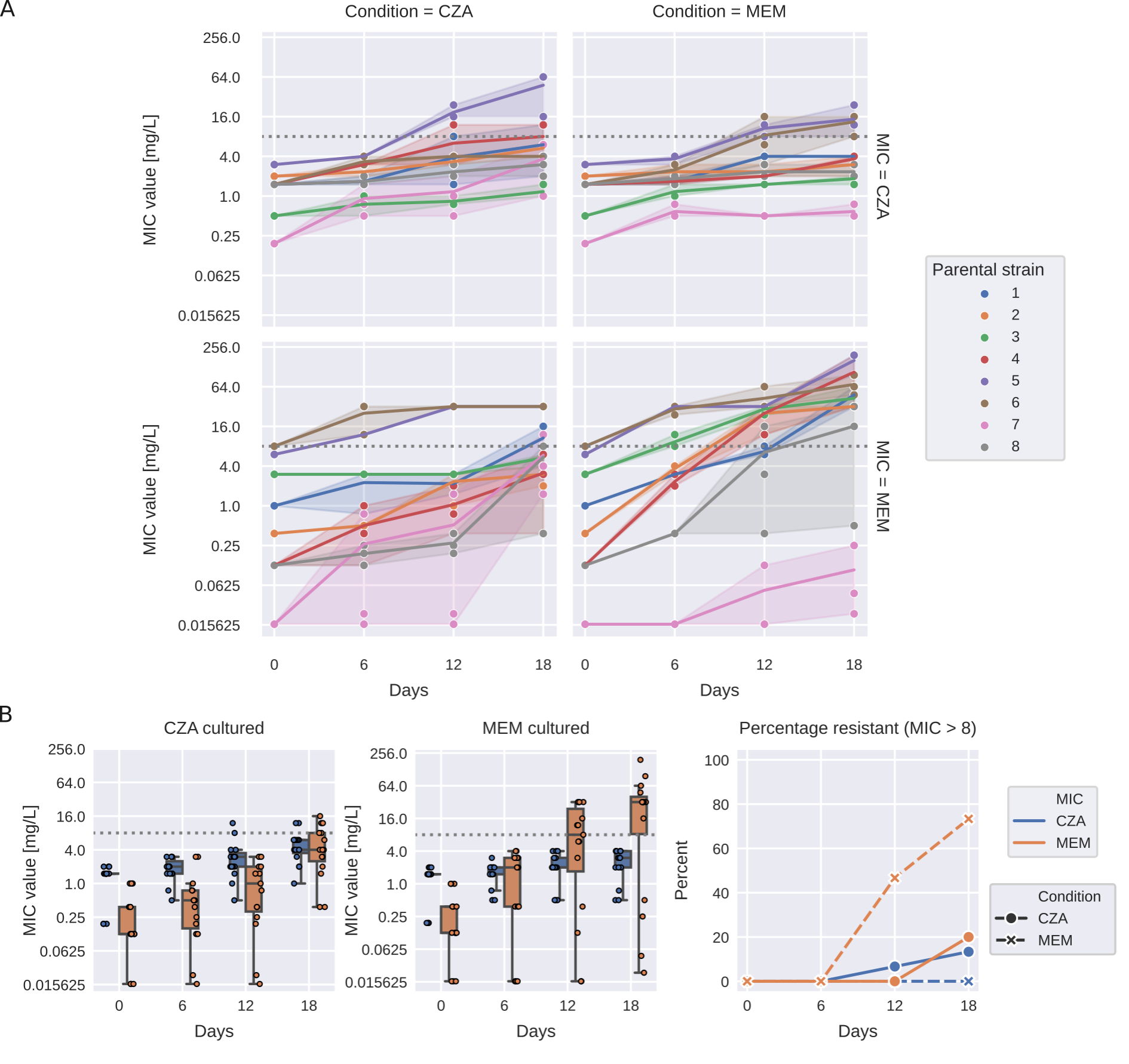

**Supplementary Figure 1**. ***Experimental evolution of antimicrobial resistance to MEM and CZA in highly sensitive clinical isolates. (****A) Resistance evolution to MEM and CZA of all strains coloured by the parental strain with the line representing the mean and shaded area representing the 95% confidence interval. Gray horizontal dotted line corresponds to the MIC=8 mg/L threshold. (B) Resistance evolution to MEM and CZA in highly sensitive parental strains (MEM MIC <=1 mg/L, n=5) exposed to CZA (left) or MEM (middle). Percentage of MEM- and CZA-resistant strains (MIC>8 mg/L) among highly sensitive parental strains exposed to either MEM or CZA (right). Gray horizontal dotted line corresponds to the MIC=8 mg/L threshold. Boxplots represent median and interquartile range (between the first and the third quartile), while the whiskers extend from the box to the farthest data point lying within 1.5x the interquartile range from the box for n=24 points (n=8 strains in triplicates). MIC - minimal inhibitory concentration.*

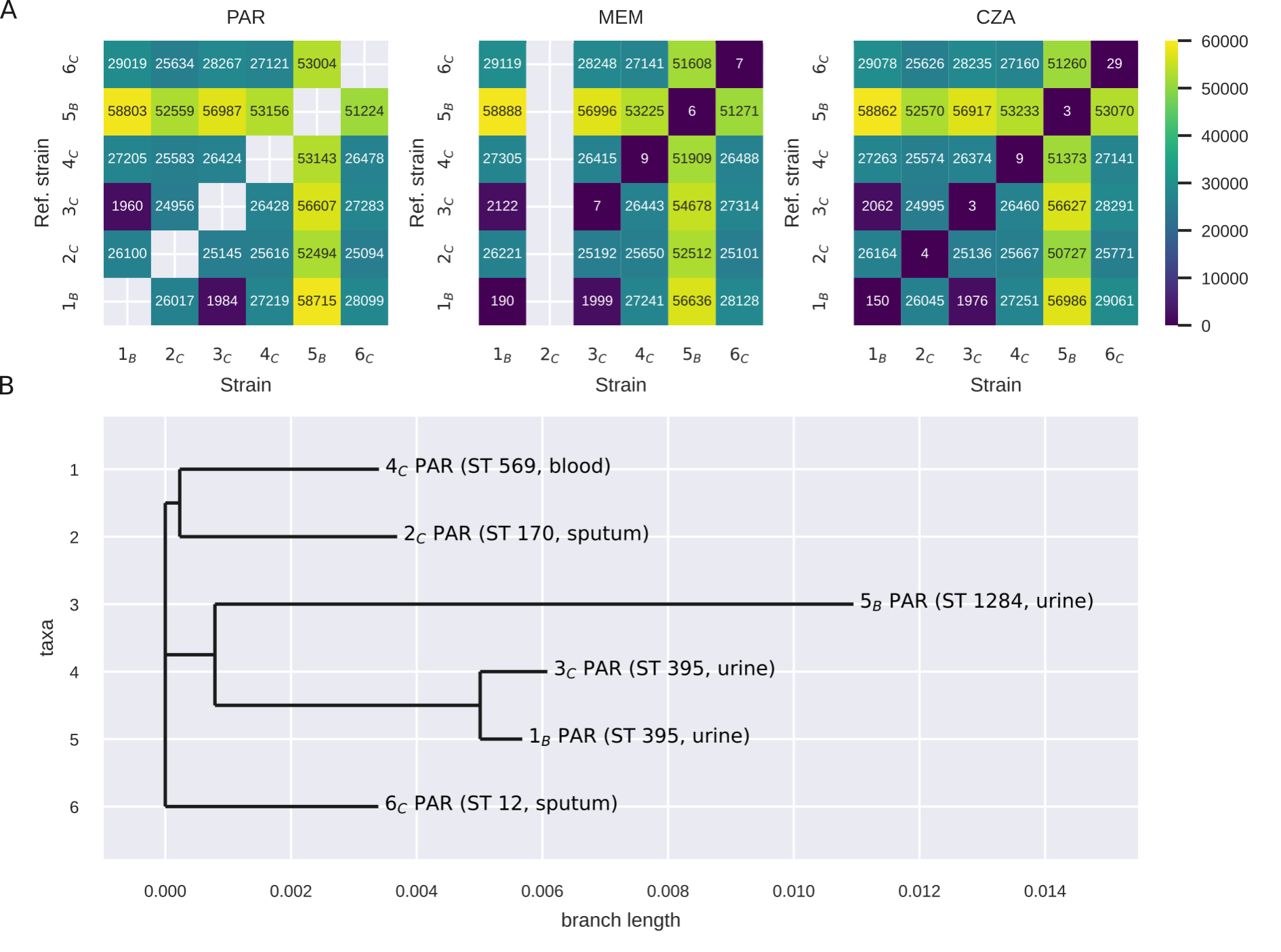

**Supplementary Figure 2**. ***Genetic diversity between the parental strains and antibiotic-evolved strains.*** *(A) Left, detected number of mutations between each pair of parental strains (P); middle, detected number of mutations between each MEM-evolved strain (x-axis) and each parental strain (y-axis); right, detected number of mutations between each CZA-evolved strain (x-axis) and each parental strain (y-axis). (B) Phylogenetic tree of parental strains with sequence types (ST) and isolation source indicated in brackets.*

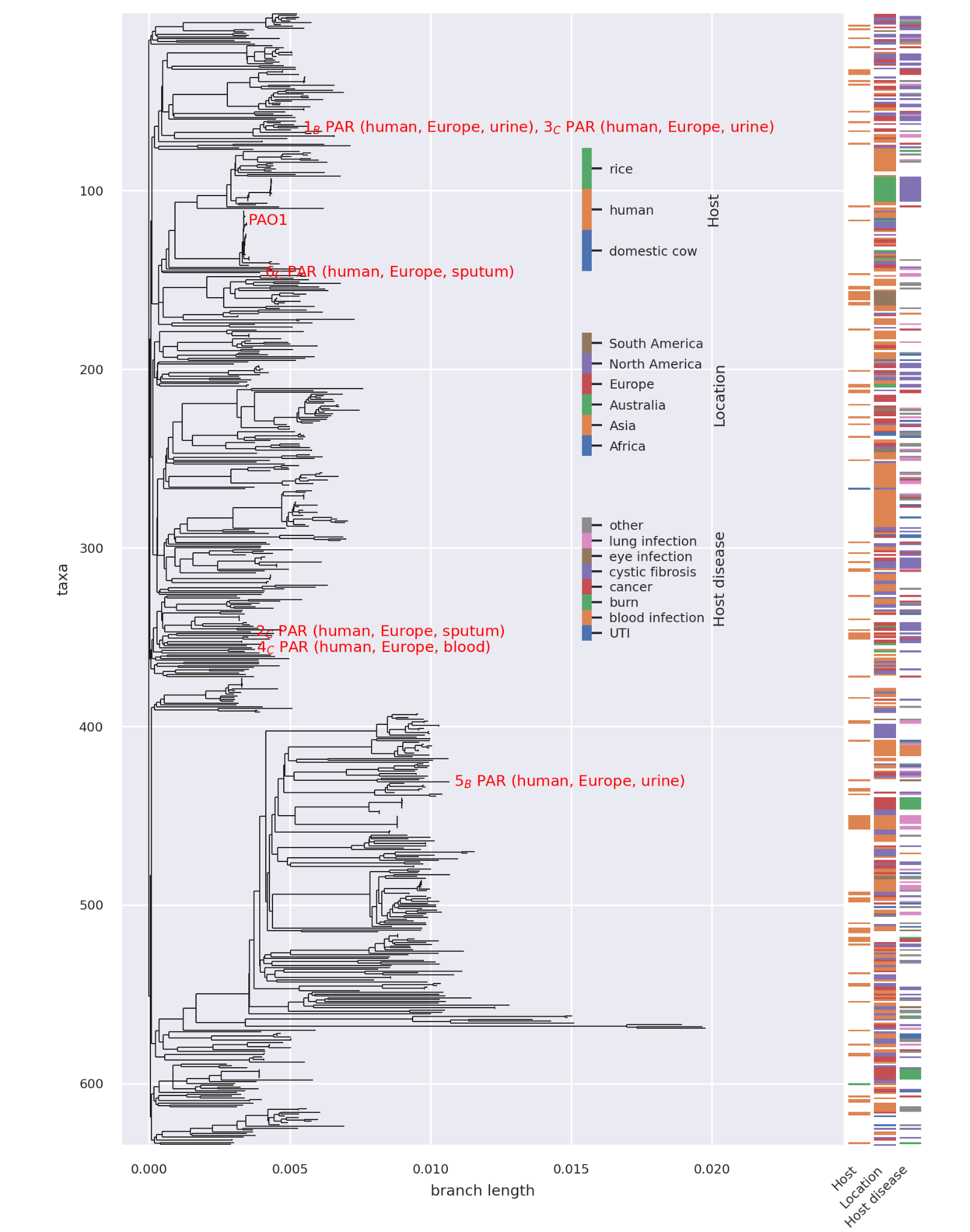

**Supplementary Figure 3**. ***Phylogenetic tree of P. aeruginosa strains with publicly available complete genomes alongside the parental strains from this study.*** *The parental strains from this study and the reference type strain PAO1 are labeled in red. Additional metadata, including the host from which each strain was collected, the geographic location of collection, and the co-occurring host disease associated with P. aeruginosa infection, are presented to the right of the phylogenetic tree. Labels of the strains in this study, highlighted in red, contain the information about the host from which each strain was collected, geographic location and source of collection.*

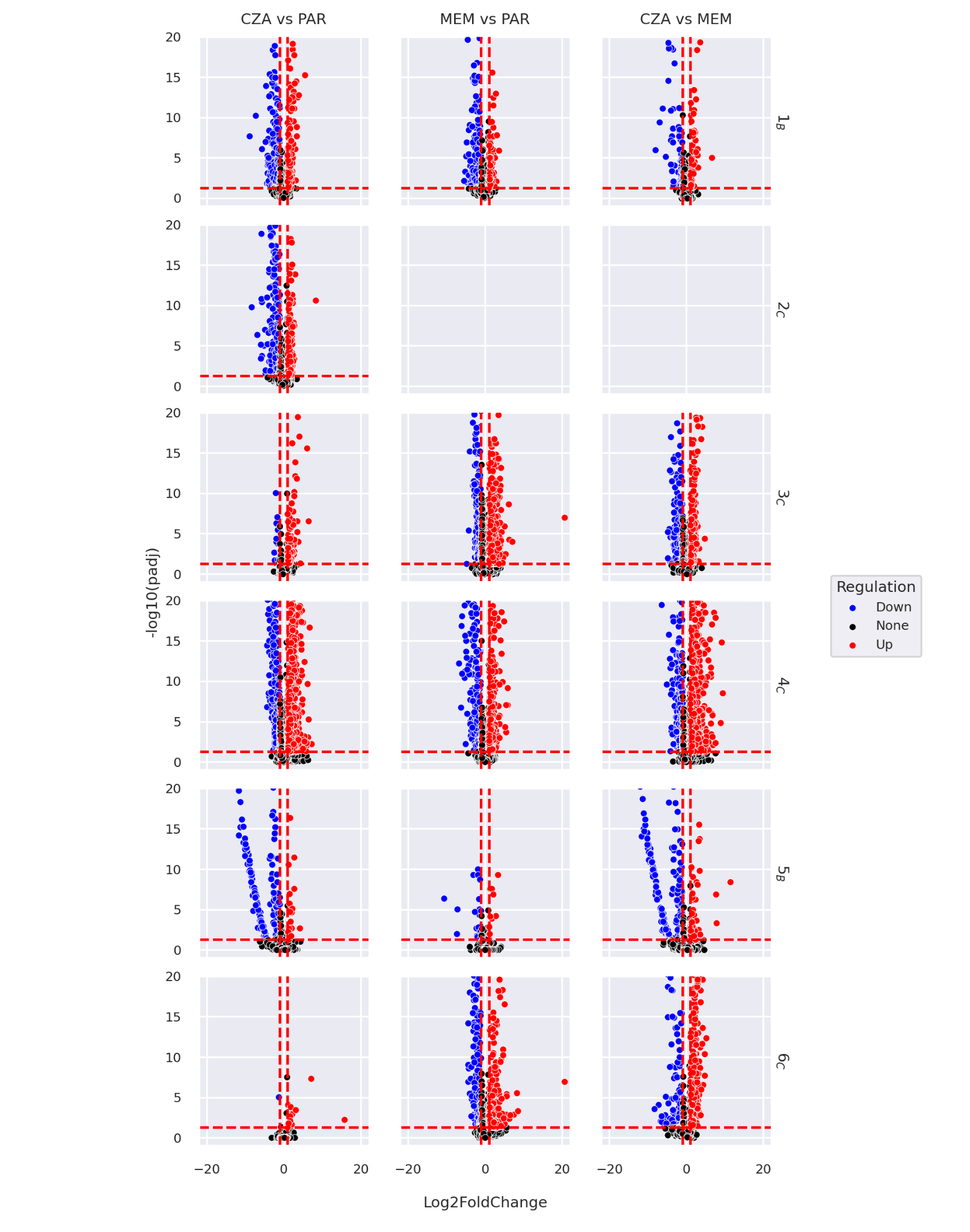

**Supplementary Figure 4. Volcano plots for transcriptomics.** Red dotted lines correspond to abs(log2FC)>1 (vertical) and padj<0.05 (horizontal) thresholds.

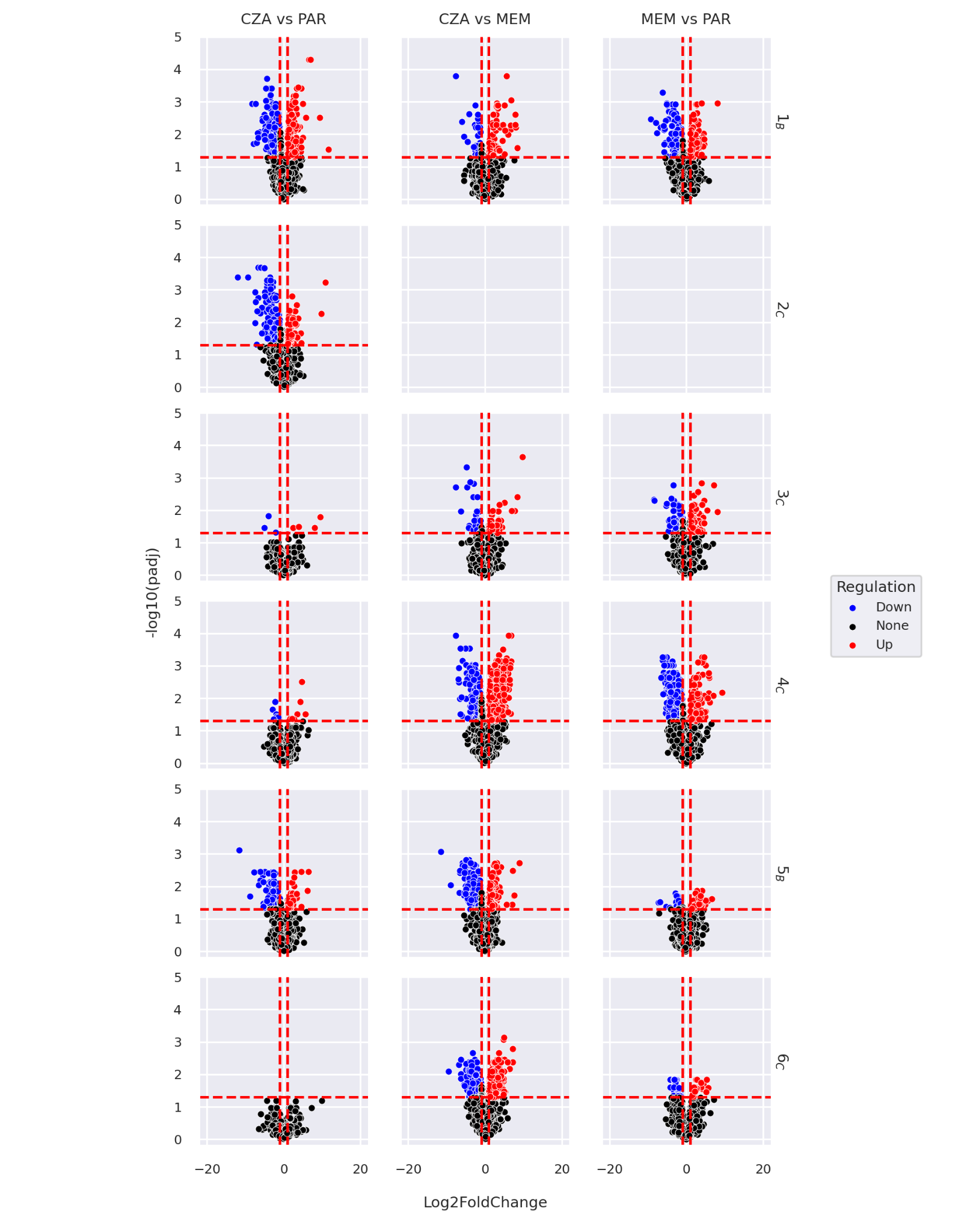

**Supplementary Figure 5. Volcano plots for proteomics.** Red dotted lines correspond to abs(log2FC)>1 (vertical) and padj<0.05 (horizontal) thresholds.

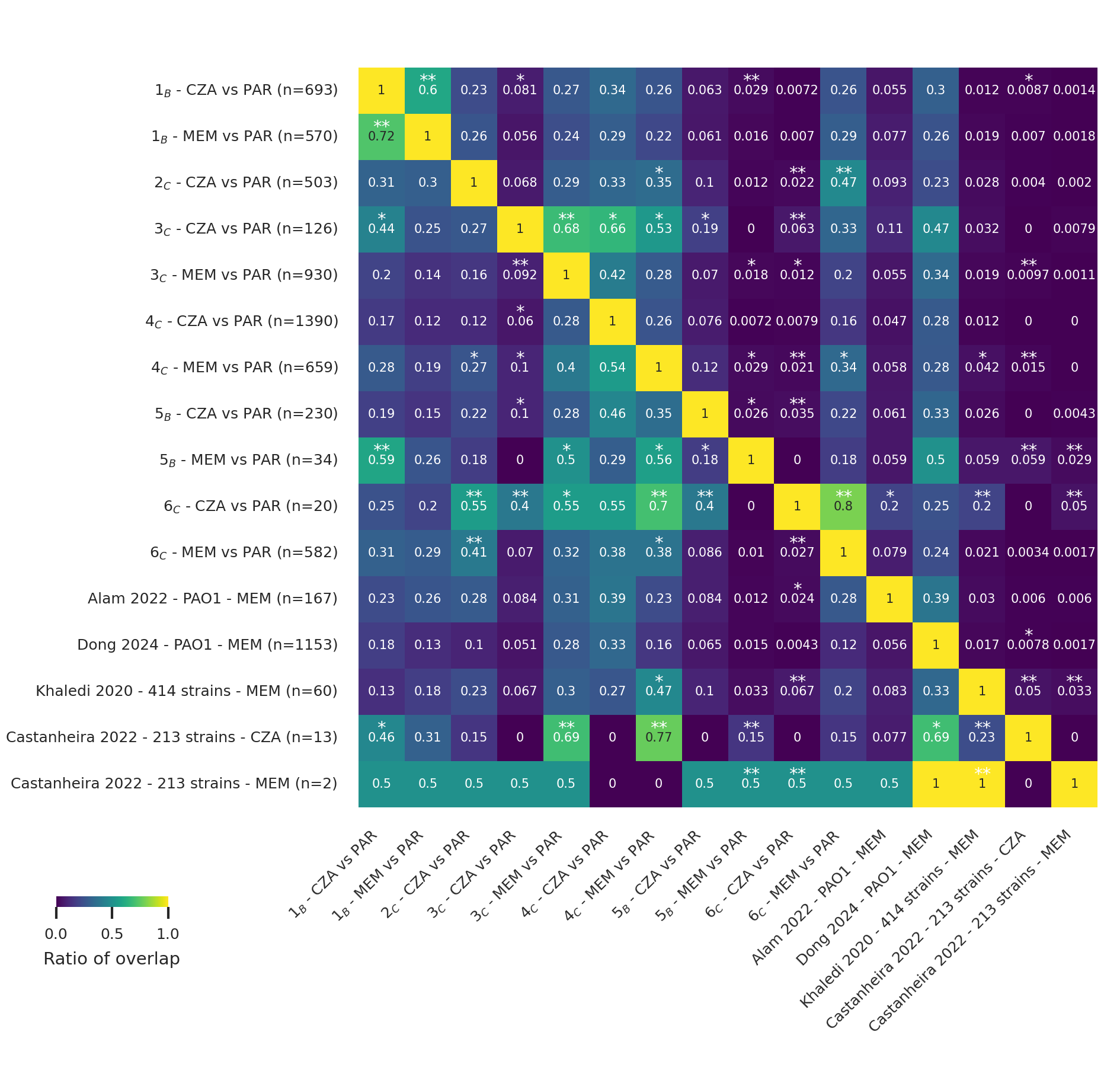

**Supplementary Figure 6**. ***Heatmap comparing significantly up- or down-regulated genes from our dataset to significantly up- or down-regulated genes from publicly available datasets.*** *The numbers in the cells represent the ratio of the number of genes in the intersection between the set of significantly up- or down-regulated genes in the row and the column to the number of genes differentially expressed in the condition indicated in the row. An asterisk in a cell indicates adjusted p-value less than 0.05 with odds ratio greater than 5 and two asterisks indicate adjusted p-value less than 0.05 with odds ratio greater than 10 calculated using Fisher’s exact test with Benjamini-Hochberg correction of p-values.*

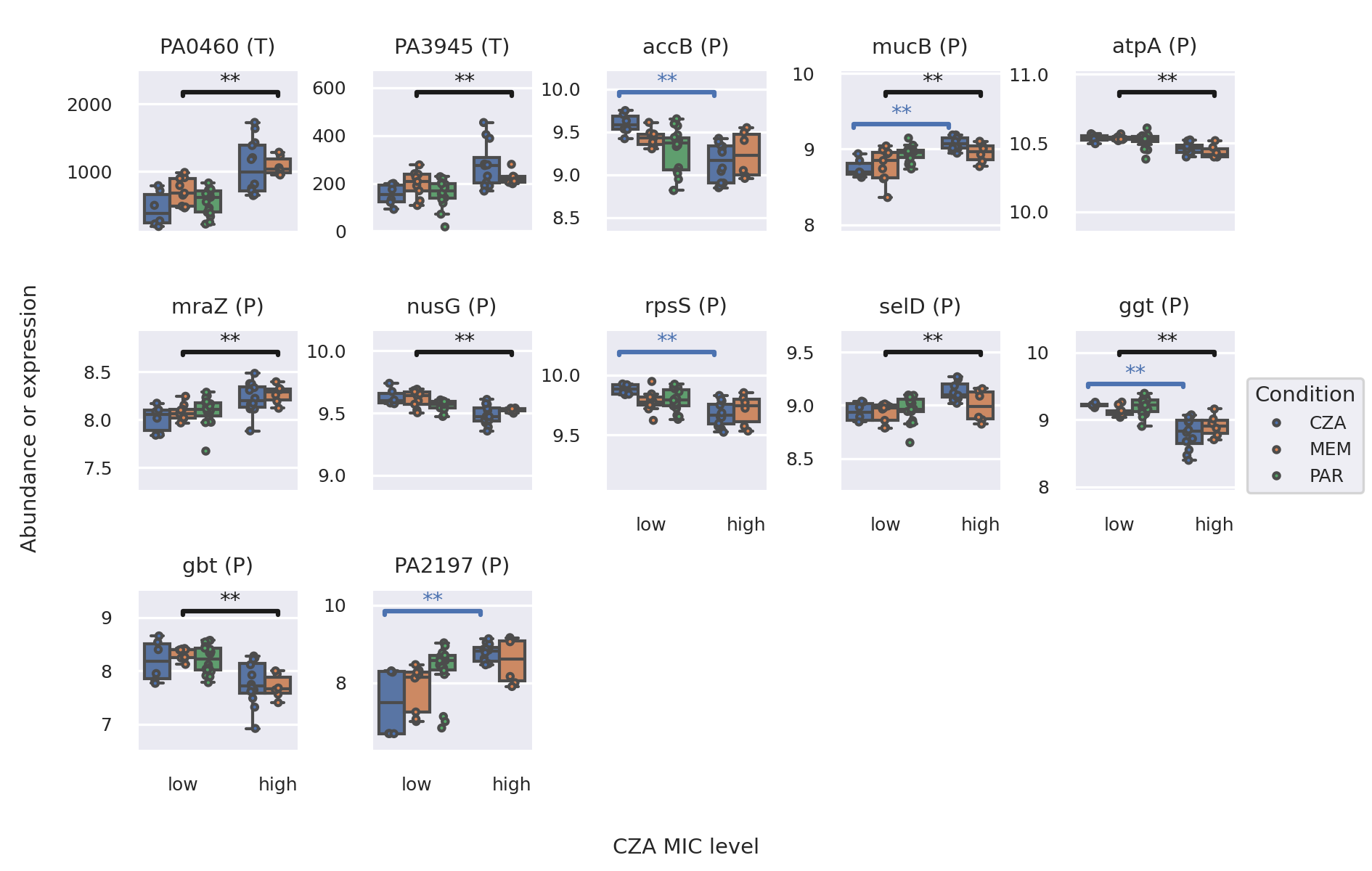

**Supplementary Figure 7**. Transcript and protein abundances of genes and proteins selected by the PLS-DA models between sensitive and resistant strains to CZA.

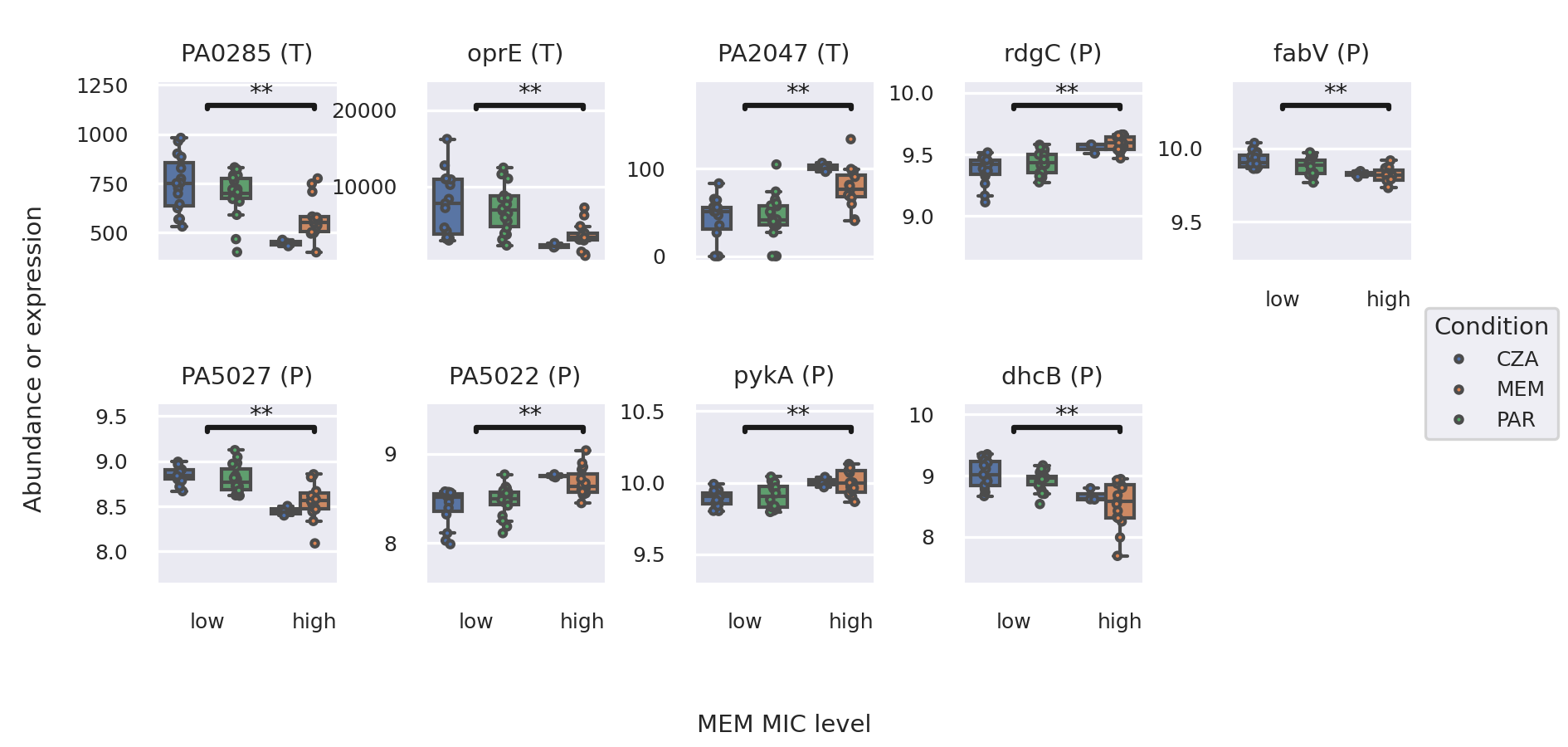

**Supplementary Figure 8**. Transcript and protein abundances of genes and proteins selected by the PLS-DA models between sensitive and resistant strains to MEM.

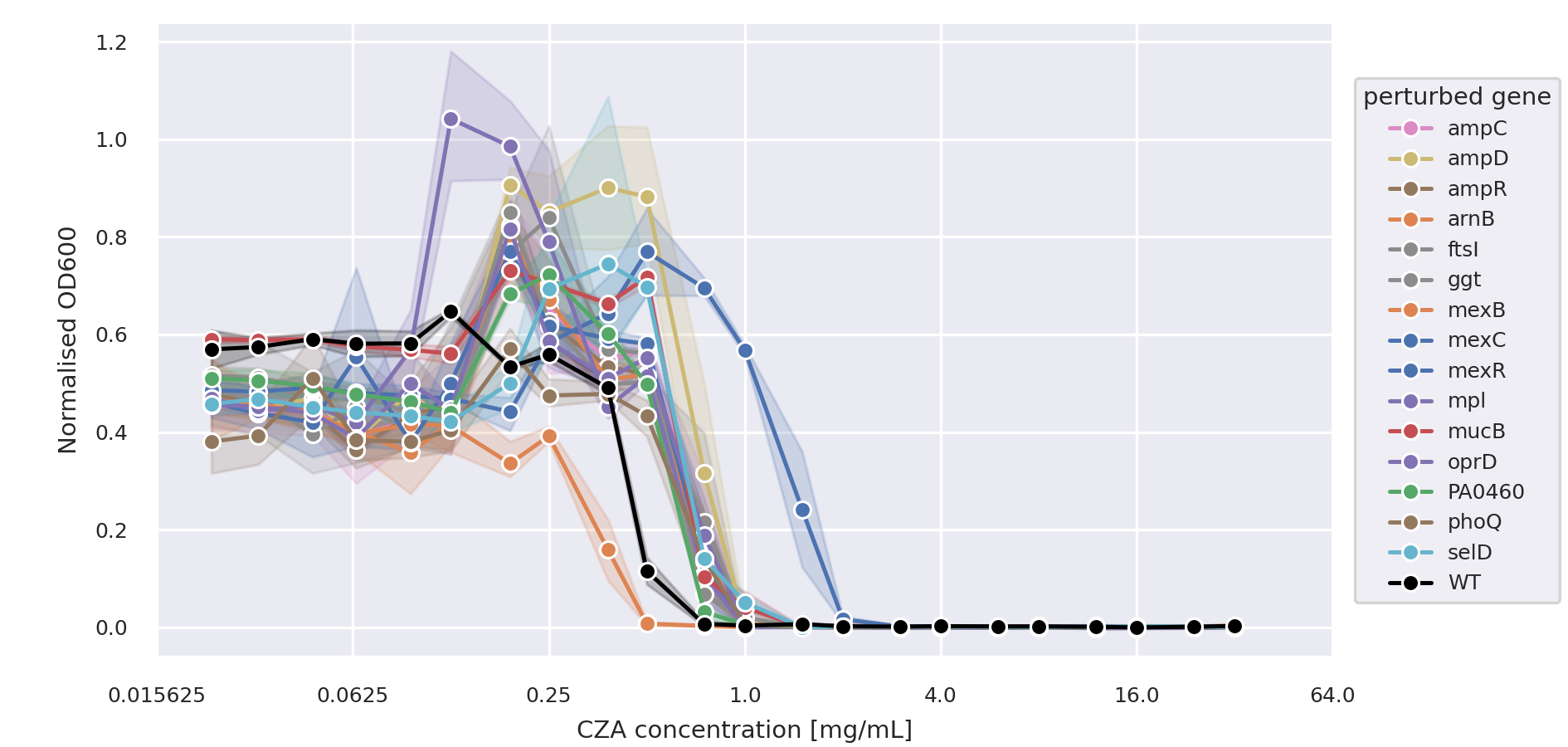

**Supplementary Figure 9**. ***MIC assays for selected transposon mutants of P. aeruginosa mPAO1.*** *Optical density (OD600) measured at each concentration of the CZA antibiotic normalised by the OD600 measurement of the blank wells. Plots represent mean and standard deviation of n=4 biological replicates.*

**Supplementary Tables**

**Supplementary Table 1.** Information about clinical isolates including the initial antibiotic resistance screening.

**Supplementary Table 2.** Evolution of resistance to MEM and CZA measured over time.

**Supplementary Table 3.** Final CZA and MEM MIC and passage length for the strains subjected to multi-omics profiling.

**Supplementary Table 4.** Trek panel results.

**Supplementary Table 5.** Mutations identified in the evolved strains with whole genome sequencing. a. List of mutations. b. Number of mutations per gene and condition. c. Number of mutations per gene.

**Supplementary Table 6.** Transcriptomic changes in the evolved strains. a. Raw gene count table. b. Normalised gene count table. c. Fold changes and p-values for all detected genes.

**Supplementary Table 7.** Proteomics changes in the evolved strains. a. Raw protein count table. b. Normalised protein count table. c. Fold changes and p-values for all detected proteins.

**Supplementary Table 8.** Gene ontology enrichment results for differentially expressed genes and proteins. a. Gene ontology enrichment results for differentially expressed genes and proteins. b. Gene ontology enrichment results for differentially expressed genes and proteins that appear in more than a given number of strains. c. Gene ontology enrichment results for genes or proteins that appeared in top hundred features in at least five PLS-DA models.

**Supplementary Table 9.** Number of strains in which each gene/protein changed upon antibiotic treatment.

**Supplementary Table 10.** Presence of significantly up- or down-regulated genes from MEM vs PAR comparison in public datasets.

**Supplementary Table 11.** Information on intersections of significantly up- or down-regulated genes between our dataset and publicly available datasets.

**Supplementary Table 12.** PLS-DA model and feature information. a. CZA and MEM resistance prediction accuracy for the full and leave-one-strain-out models. b. Model feature importance. c. The count of models in which each gene ranked among the top 100 features based on weights. d. Gene ontology enrichment results for features selected by a given number of models.

**Supplementary Table 13.** Fold change and p-values of gene expression and protein abundances between all strains grouped by resistance to CZA or MEM across conditions.

**Supplementary Table 14.** Top genes and proteins selected from PLS-DA analysis.

**Supplementary Table 15**. Top genes and proteins selected from mutation analysis, differential analysis, and PLS-DA.

**Supplementary Table 16.** Transposon mutant MIC screening results. a. Raw and normalised OD600 values. b. MIC value for each mutant.

**Supplementary Table 17.** Primers designed for the CRISPR-Cas9 editing.

**Supplementary Table 17. Primers designed for the CRISPR-Cas9 editing.**

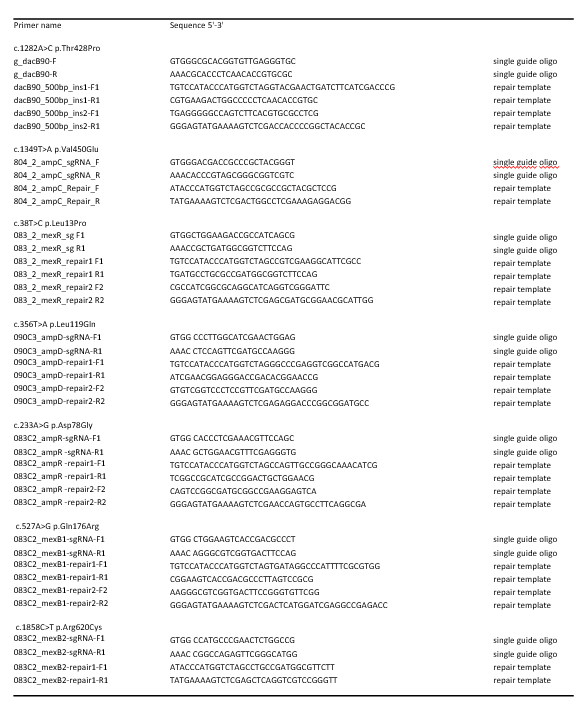
